# Yorkie-independent negative feedback couples Hippo pathway activation with Kibra degradation

**DOI:** 10.1101/2020.07.08.192765

**Authors:** Sherzod A. Tokamov, Ting Su, Anne Ullyot, Richard G. Fehon

## Abstract

The Hippo signaling pathway regulates tissue growth in many animals. Multiple upstream components are known to promote Hippo pathway activity, but the organization of these different inputs, the degree of crosstalk between them, and whether they are regulated in a distinct manner is not well understood. Kibra activates the Hippo pathway by recruiting the core Hippo kinase cassette to the apical cortex. Here we show that the Hippo pathway downregulates Kibra levels independently of Yorkie-mediated transcriptional output. We find that the Hippo pathway promotes Kibra degradation via SCF^Slimb^-mediated ubiquitination, that this effect requires the core kinases Hippo and Warts, and that this mechanism functions independently of other upstream Hippo pathway activators including Crumbs and Expanded. Moreover, Kibra degradation appears patterned across tissue. We propose that Kibra degradation by the Hippo pathway serves as a negative feedback loop to tightly control Kibra-mediated Hippo pathway activation and ensure optimally scaled and patterned tissue growth.

## Introduction

How organs achieve and maintain optimal size is a fundamental question in developmental biology. The Hippo signaling pathway is an evolutionarily conserved inhibitor of tissue growth that was first identified in *Drosophila* in somatic mosaic screens for tumor-suppressor genes (Xu et al., 1995; Tapon et al., 2002; Harvey et al., 2003; Wu et al., 2003). Central to the Hippo pathway activity is a kinase cassette that includes serine/threonine kinases Tao-1, Hippo (Hpo), and Warts (Wts), as well as two scaffolding proteins Salvador (Sav) and <u>M</u>ob <u>a</u>s <u>t</u>umor <u>s</u>uppressor (Mats). Activation of the Hippo pathway results in a kinase cascade that culminates in the phosphorylation of a transcriptional co-activator Yorkie (Yki) by Wts, which inhibits Yki nuclear accumulation. Conversely, inactivation of the Hippo pathway allows Yki to translocate into the nucleus where, together with its DNA-binding partners such as Scalloped (Sd), it promotes transcription of pro-growth genes. As a result, inactivation of the Hippo pathway is characterized by excessive tissue growth. Mutations that disrupt Hippo pathway activity can lead to various human disorders including benign tumors and carcinomas (Zheng and Pan, 2019).

A distinct feature of the Hippo pathway is the remarkably complex organization of its upstream regulatory modules (Fulford et al., 2018). The core kinase cascade is regulated from the junctional and apical cortex of epithelial cells by multiple upstream components, including Fat (Ft), Dachsous (Ds), Echinoid (Ed), Expanded (Ex), Crumbs (Crb), Kibra (Kib) and Merlin (Mer). Ft and Ds are protocadherins that promote Hippo pathway activity by restricting the activity of Dachs, an atypical myosin that inhibits Wts (Bennett and Harvey, 2006; Cho et al., 2006; Mao, 2006; Matakatsu and Blair, 2012; Vrabioiu and Struhl, 2015). Ed is a cell-cell adhesion protein that binds and stabilizes Sav at the junctional cortex, thereby enabling Sav to promote Hippo pathway activity (Yue et al., 2012). Ex is a FERM-domain protein that also localizes at the junctional cortex where it binds to the transmembrane protein Crb and activates the Hippo pathway by recruiting the core kinase cassette (Hamaratoglu et al., 2006; Ling et al., 2010; Robinson et al., 2010; Sun et al., 2015). The WW-domain protein Kib and FERM-domain protein Mer localize both at the junctional and apical medial cortex and promote Hippo pathway activity by recruiting the core kinase cassette independently of Ex (Yu et al., 2010; Baumgartner et al., 2010; Genevet et al., 2010; Hamaratoglu et al., 2006; Su et al., 2017). The existence of multiple upstream regulatory modules that converge to control the activity of a single downstream effector, Yki, raises a question of whether and how these parallel inputs are regulated and to what extent they are distinct from one another.

One way that cells modulate signaling output is by controlling the levels of signaling components. Within the Hippo pathway, the transcription of Ex, Kib, and Mer is positively regulated by Yki activity to form a negative feedback loop (Hamaratoglu et al., 2006; Genevet et al., 2010; Yee et al., 2019). Multiple Hippo pathway components are also regulated post-translationally. For example, Crb promotes Ex ubiquitination via Skip/Cullin/F-box^Slimb^ (SCF^Slimb^) E3 ubiquitin ligase complex, which leads to Ex degradation (Ribeiro et al., 2014; Fulford et al., 2019). Similarly, Ds and Dachs levels are downregulated by the SCF^Fbxl-7^ E3 ubiquitin ligase, and Dachs stability is also influenced by an E3 ubiquitin ligase called Early girl (Rodrigues-Campos and Thompson, 2014; Misra and Irvine, 2019). Sav stability is also inhibited by the HECT ubiquitin ligase Herc4 (Aerne et al., 2015). These studies underscore the importance of post-translational regulation of Hippo pathway components and suggest that individual signaling branches of the Hippo pathway might be regulated in a distinct manner from one another.

In this study, we reveal that Hippo pathway activity negatively regulates Kib levels via a post-translational negative feedback. We show that the regulation of Kib levels by the Hippo pathway is independent of Yki- and Sd-mediated transcriptional output and is instead mediated by SCF^Slimb^. We find that this mechanism operates independently of other upstream inputs such as Ex/Crb, Ft/Ds, or Ed. Intriguingly, our data suggest that Kib degradation is patterned across the wing imaginal tissue. We propose a model in which Kib-mediated Hippo pathway activation results in Kib degradation in isolation from other upstream inputs, thereby forming a tightly compartmentalized negative feedback loop. Such feedback may function as a homeostatic mechanism to tightly control signaling output specifically downstream of Kib and ensure proper tissue growth during development.

## Results

### Transcriptional feedback is insufficient to explain the increase in Kibra abundance upon pathway inactivation

A notable feature of the Hippo pathway is that its upstream components Kib, Ex, and Mer are upregulated by Yki transcriptional activity in a negative feedback loop (Hamaratoglu et al., 2006; Genevet et al., 2010; Yee et al., 2019). In particular, Kib levels were previously shown to be significantly elevated in double-mutant *Mer; ex* somatic mosaic clones, consistent with the transcriptional feedback regulation of *kibra* by Yki (Genevet et al., 2010). However, when we examined endogenous Kib::GFP in live wing imaginal discs containing either *Mer* or *ex* mutant clones individually, we found that Kib abundance was significantly higher in *Mer* mutant clones than in *ex* mutant clones (Figures 1A-1B’, quantified in Figure 1C). These results suggest that loss of Mer has a greater effect on Yki transcriptional activity than loss of Ex, which has not been reported previously.

**Figure 1.**
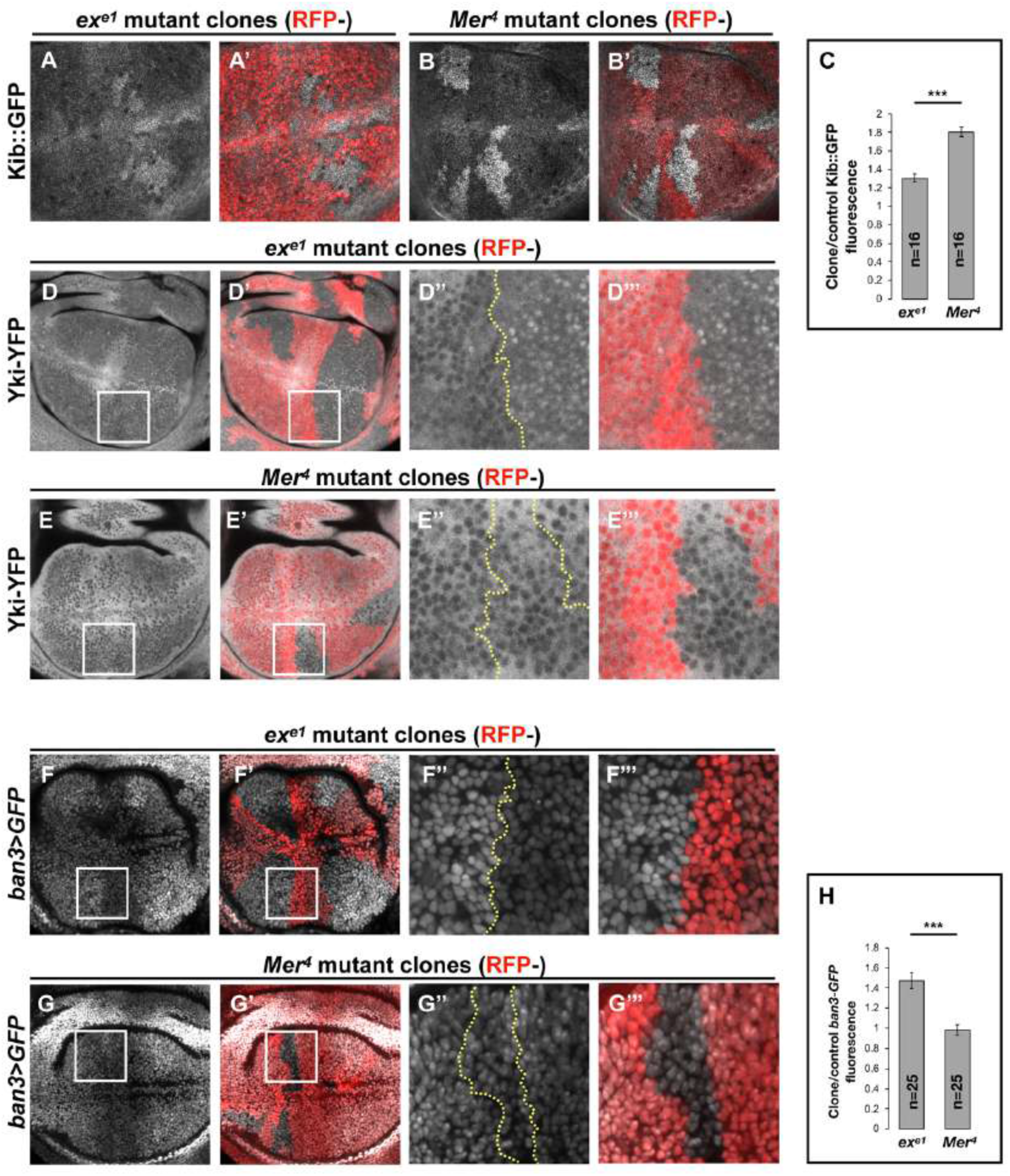
Transcriptional feedback alone does not explain Kibra upregulation in *Mer* clones. (A-G”’) All tissues shown are living late third instar wing imaginal discs expressing the indicated fluorescent proteins. (A-C) Endogenous Kib::GFP in *ex* (A and A’) or *Mer* (B and B’) somatic mosaic clones (indicated by loss of RFP). Loss of Mer leads to a greater increase in Kib levels than loss of Ex. Quantification is shown in (C). (D-E”’) Endogenously expressed Yki-YFP is strongly nuclear in *ex* mutant clones (DD”’) but is mostly cytoplasmic in *Mer* mutant clones (E-E”’). (F-H) Expression of a Yki activity reporter, *ban3>GFP*, is elevated in *ex* mutant clones (F-F”’) but is not detectably affected in *Mer* mutant clones (G-G”’). Quantification is shown in (H). Yellow dashed line indicates clone boundary. Quantification in C & H is represented as the mean ± SEM; n = number of clones; \*\*\**p* ≤ 0.001.

To directly assess the relative contribution of Mer and Ex to Yki activity, we examined the nuclear localization of endogenously expressed Yki-YFP, a biosensor for Yki activity (Su et al., 2017; Xu et al., 2018). In sharp contrast to what we observed with Kib levels, Yki strongly accumulated in the nuclei of *ex* mutant clones, whereas Yki was mostly cytoplasmic and indistinguishable from wild-type cells in *Mer* clones (Figures 1D-1E”’). These results indicate that Ex is more potent at inhibiting Yki nuclear translocation than Mer, consistent with Ex’s ability to limit Yki activity by direct sequestration at the junctional cortex (Badouel et al., 2009) and suggesting that loss of *ex* should have a greater effect on pathway target gene expression than loss of *Mer.*

To compare the effects of *Mer* and *ex* loss on target gene expression, we examined the expression of *ban3>GFP* (Matakatsu and Blair, 2012), a reporter for one of Yki’s target genes *bantam* (Thompson and Cohen, 2006; Nolo et al., 2006). *ban3>GFP* expression was significantly upregulated in *ex* mutant clones (Figures 1F-1F’”), whereas no detectible difference was observed in *Mer* clones relative to control tissue (Figures 1G-1G”’, quantified in Figure 1H), indicating that Yki is more active in *ex* clones than in *Mer* clones. Together, these results suggest that the dramatic increase in Kib levels in *Mer* clones cannot be explained strictly by Yki-mediated transcriptional feedback and suggest that Kib is also regulated via a previously unrecognized non-transcriptional mechanism.

### Hippo pathway activity regulates Kibra abundance non-transcriptionally

If a Yki-independent mechanism is responsible for Kib upregulation in *Mer* clones, then Kib levels should be elevated in *Mer* clones in the absence of Yki activity. To test this hypothesis, we took advantage of a previously published method of blocking Yki-mediated transcription downstream of the Hippo pathway by removing Yki’s DNA binding partner, Sd, in the eye imaginal disc, where Sd is dispensable for cell viability (Koontz et al., 2013; Yu and Pan, 2018). Endogenous Kib::GFP was upregulated in *sd; Mer* double mutant clones to a similar degree as in *Mer* single mutant clones (Figures S1A-S1B’), suggesting that Mer regulates Kib levels independently of Yki activity.

To understand how Mer regulates Kib levels, we set out to develop a simpler approach to uncouple Kib protein abundance from its transcriptional regulation. Recently, the *ubiquitin 63E* promoter was used to drive expression of other Hippo pathway components to study their post-translational regulation (Aerne et al., 2015; Fulford et al., 2019), based on the assumption that the ubiquitin promoter is not regulated by Yki activity. Therefore, we made a transgenic fly line ectopically expressing Kibra-GFP-FLAG under control of the ubiquitin promoter (Ubi>Kib-GFP) (Fig. 2A). Similar to endogenous Kib::GFP, Ubi>Kib-GFP localized both at the junctional and medial cortex (Figure S1D). Additionally, flies expressing Ubi>Kib-GFP had slightly undergrown wings compared to control flies expressing Ubi>GFP (Figure S1E), suggesting that Ubi>Kib-GFP promotes Hippo pathway activity.

**Figure 2.**
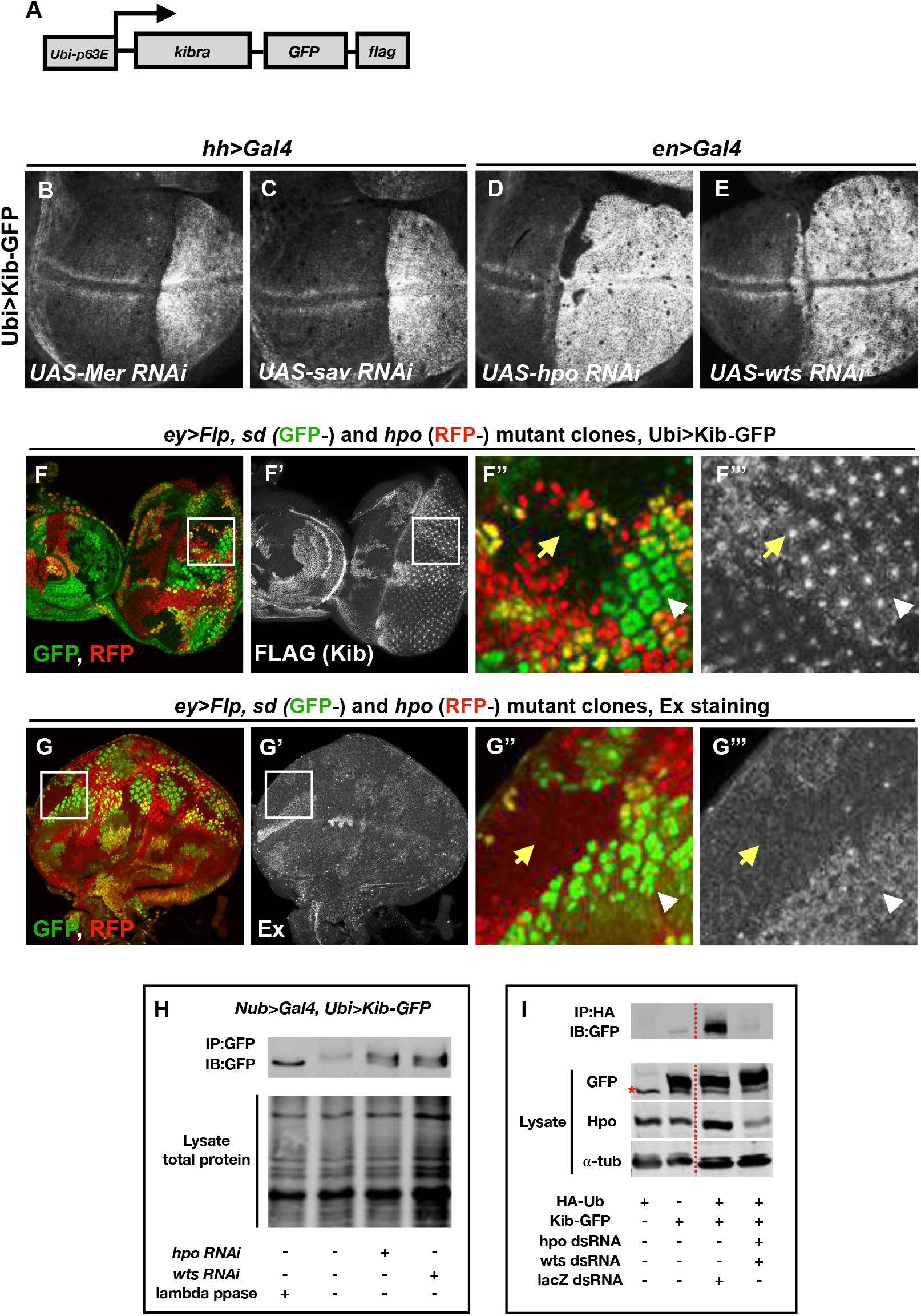
The Hippo pathway regulates Kibra levels independent of Yki-mediated transcription. A) A cartoon of the DNA construct used to generate the Ubi>Kib-GFP transgenic fly line. B-E) Depletion of Hippo pathway components Mer, Sav, Hpo and Wts by RNAi in the posterior compartment of the wing results in elevated Ubi>Kib-GFP levels. F-F”’) In the eye imaginal disc, Ubi>Kib-GFP is upregulated both in *hpo* mutant clones and *sd; hpo* double mutant clones, indicating that Hippo pathway activity controls Kib levels independently of Yki/Sd-mediated transcription. White arrowheads indicate *hpo* single mutant clones; yellow arrows indicate *sd; hpo* double mutant clones. See also Figure S2E. G-G”’) Ex levels are also upregulated in *hpo* mutant clones; but in contrast to Kib, Ex upregulation is not observed in *sd; hpo* double mutant clones. H) Kib is phosphorylated in wing discs, and depletion of Hpo or Wts leads to decreased Kib phosphorylation. I) Kib is ubiquitinated in S2 cells, and depletion of Hpo and Wts leads to decreased Kib ubiquitination. Asterisk indicates a non-specific band. The dashed line indicates that a lane was cropped out.

Consistent with the hypothesis that Mer negatively regulates Kib levels non-transcriptionally, depletion of Mer in the posterior compartment of the wing disc using the *hh>Gal4* driver led to a substantial increase in Ubi>Kib-GFP levels (Figure 2B). Knockdown of Sav, Hpo, and Wts also dramatically increased Ubi>Kib-GFP levels (Figures 2C–2E), suggesting that Hippo pathway activity negatively regulates Kib abundance. In contrast, expression of a Ubi>RFP control transgene was not affected by depletion of Hpo, confirming that Yki does not regulate expression at the ubiquitin promoter (Figures S1C-S1C”). Ubi>Kib-GFP appeared to accumulate junctionally as we depleted the downstream pathway components Hpo and Wts, suggesting that Hippo pathway activity might regulate both Kib abundance and its localization at the junctional cortex (Figure S1F).

Ex is also upregulated upon Hippo pathway inactivation, with a particularly strong increase when Hpo or Wts is depleted (Hamaratoglu et al., 2006, and Figures S2A-S2D). Ex and Kib also form a complex in cultured cells (Genevet et al., 2010; Yu et al., 2010), raising the possibility that the increase in Ubi>Kib-GFP levels upon Hpo or Wts depletion is caused by increased interaction with Ex resulting in greater Kib stability. To test this possibility, we compared Ex and Ubi>Kib-GFP levels in *hpo* or *sd; hpo* double mutant clones. While Ubi>Kib-GFP levels were similarly elevated in both *hpo* and *sd; hpo* double mutant clones (Figures 2F–2F”’, Figure S2E), Ex levels were upregulated only in *hpo* single mutant clones but not in *sd; hpo* double mutant clones (Fig. 2G-G”’), indicating that the increase in Kib levels upon Hippo pathway inactivation is not mediated via Ex. Furthermore, transient co-depletion of Hpo and Yki in the wing disc posterior compartment using Gal80^ts^ did not suppress the increase in Kib abundance observed when Hpo alone was depleted, even though Yki was sufficiently depleted to suppress tissue overgrowth induced by Hpo depletion alone (Figure S2F). Together, these results provide strong evidence that the Hippo pathway regulates Kib levels independently of Yki transcriptional output.

### The Hippo pathway promotes Kibra phosphorylation and ubiquitination

Our observation that Hippo pathway activity controls Kib levels in a Yki-independent manner suggests that Kib could be regulated post-translationally. Protein abundance is commonly regulated by phosphorylation-dependent ubiquitination, and multiple Hippo pathway components are regulated via ubiquitin-mediated proteasomal degradation (Ribeiro et al., 2014; Rodrigues-Campos and Thompson, 2014; Cao et al., 2014; Aerne et al., 2015; Ma et al., 2018; Ly et al., 2019). Since a kinase cascade is at the core of the Hippo pathway, we hypothesized that Hippo pathway activity could promote Kib phosphorylation and target it for ubiquitination and subsequent degradation.

We first asked whether Kib is phosphorylated in a pathway dependent manner *in vivo.* To this end, we examined the phosphorylation state of Kib-GFP in wing imaginal discs depleted for either Hpo or Wts using a gel shift assay. In wild-type controls, phosphatase treatment of immunoprecipitated Kib-GFP resulted in increased mobility and coalescence into a single band, suggesting that Kib is normally phosphorylated (Figure 2H). Depletion of either Hpo or Wts resulted in a faster migrating Kib band that aligned with phosphatase-treated Kib (Figure 2H), suggesting that Hippo pathway activity promotes Kib phosphorylation.

Next, we asked if Kib is ubiquitinated and, if so, whether this depends on Hippo pathway activity. To address this question, we expressed Kib-GFP and HA-tagged ubiquitin in cultured Drosophila S2 cells. We found that Kib was ubiquitinated (Figure 2I) and that co-depletion of the core pathway kinases Hpo and Wts resulted in dramatically decreased Kib ubiquitination (Figure 2I). Together, these results suggest that the Hippo pathway regulates Kib levels via phosphorylation-dependent ubiquitination and subsequent degradation.

### Slimb regulates Kibra levels via a consensus degron motif

To better understand how the Hippo pathway controls Kib levels via ubiquitination, we sought to identify the machinery that mediates this process. Protein ubiquitination occurs via an enzymatic cascade that culminates in the covalent attachment of ubiquitin molecules to substrates by E3 ubiquitin ligases (Zheng and Shabek, 2017). We first tested the effects of RNAi depletion or overexpression of E3 ubiquitin ligases previously reported to act within the Hippo pathway on Ubi>Kib-GFP abundance. Of these, only depletion of the F-box protein Slimb, and its partners SkpA and Cul1, increased Ubi>Kib-GFP levels (Figure 3A; Figures S3A-D). Importantly, increased Ubi>Kib-GFP was evident throughout the affected cells in comparison to control tissue (Figure 3A’), suggesting that overall Kib abundance was increased. Because Slimb also regulates Ex turnover (Ribeiro et al., 2014), we considered that increased Ex could indirectly influence Kib stability. However, co-depletion of Ex and Slimb did not suppress the increase in Ubi>Kib-GFP levels (Figures S3E-G’), suggesting that Slimb directly regulates Kib abundance.

**Figure 3:**
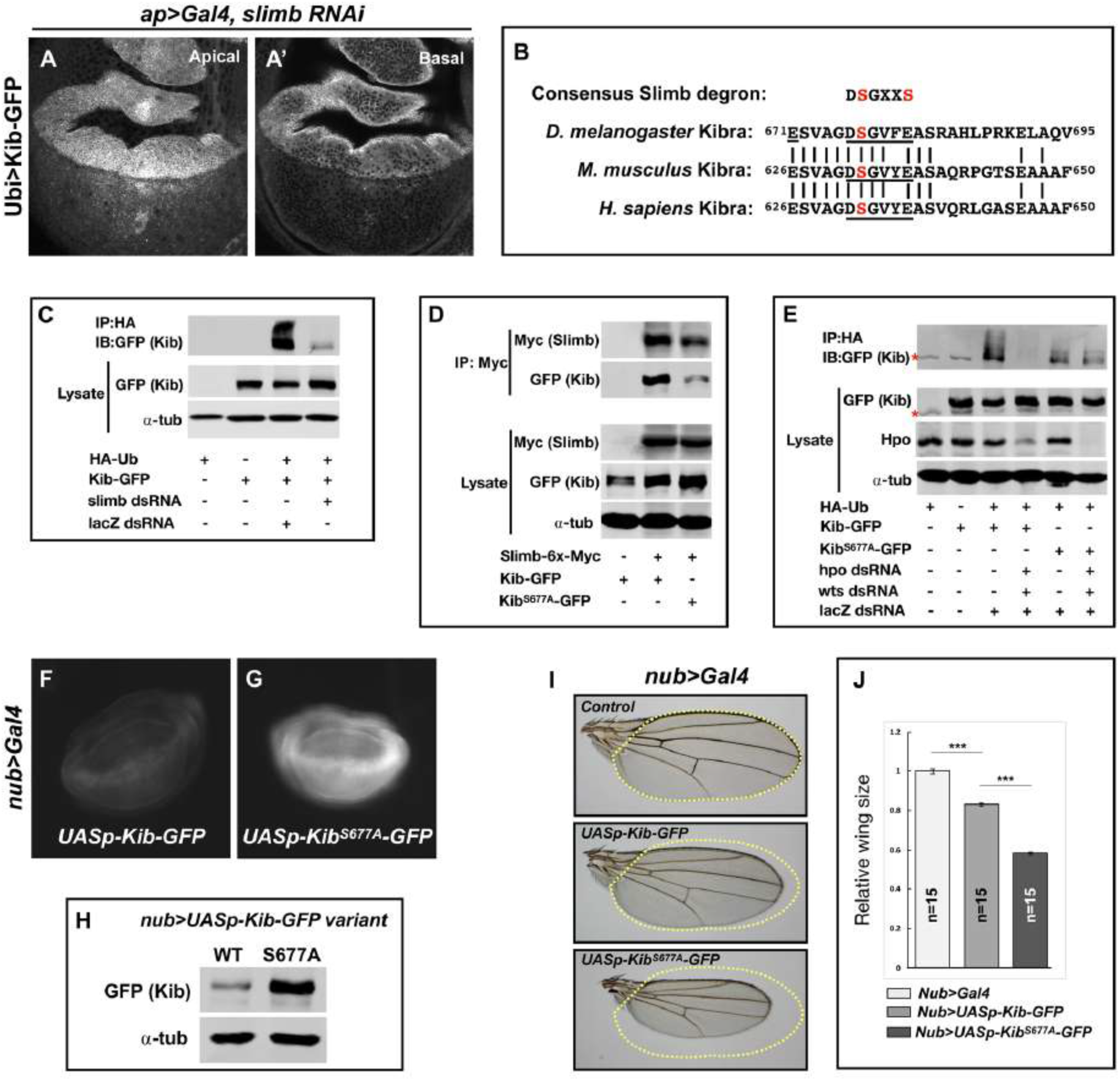
Slimb regulates Kibra abundance via a consensus degron. A-A’) Depletion of Slimb in the dorsal compartment of the wing imaginal disc results in increased Ubi>Kib-GFP levels both apically and basally. See also Figures S3A-C. B) Alignment of the fly, mouse, and human Kib protein sequences showing the conservation of the putative Slimb degron motif DSGXXS (underlined). C) Immunoblot showing that depletion of Slimb in S2 cells decreases Kib ubiquitination. Asterisk indicates a non-specific band. D) Co-IP experiments showing that Kib forms a complex with Slimb in S2 cell lysates in a degron-dependent manner. E) Ubiquitination of the degron mutant, Kib^S677A^, is diminished and is insensitive to Hippo pathway inactivation. F-G) Widefield fluorescence images of wing discs expressing either UASp-Kib-GFP (F) or UASp-Kib^S677A^-GFP (G) with the *nub>Gal4* driver; images were taken using identical settings. H) Immunoblot of wing disc cell lysates (20 discs each) of UASp-Kib-GFP or UASp-Kib^S677A^-GFP expressed with the *nub>Gal4* driver. I-J) Ectopic expression of Kib^S677A^-GFP in the wing results in stronger growth suppression than expression of wild-type Kib-GFP. Quantification of wing sizes in (I) is represented as mean ± SEM relative to the control; n = number of wings; \*\*\**p* ≤ 0.001.

Slimb is a homolog of the mammalian ß-TrCP that functions as a substrate-targeting component of the SCF E3 ubiquitin ligase complex by recognizing a consensus degron motif on target proteins (Skaar et al., 2013). Kib contains a conserved single stretch of amino acids ^676^DSGVFE^681^ that matches the consensus Slimb degron (Figure 3B). If Slimb regulates Kib stability via the degron, then we predict that 1) Kib ubiquitination should be Slimb-dependent, 2) Slimb should physically interact with Kib via the degron, 3) mutation of the degron site should diminish Kib ubiquitination, and 4) the degron mutant Kib should display greater stability than wildtype Kib. Using S2 cells, we found that depletion of Slimb severely reduces Kib ubiquitination and that Kib and Slimb formed a complex (Figures 3C and 3D). Additionally, mutating a serine residue in Kib (Kib^S677A^) known to be important for proper substrate recognition by Slimb (Hart et al., 1999; Rogers et al., 2009; Morais-de-Sá et al., 2013; Ribeiro et al., 2014) significantly reduced both Slimb-Kib interaction (Figure 3D) and Kib ubiquitination (Figure 3E).

To assess the effects of the degron mutation on protein stability *in vivo*, we generated wild-type and Kib^S677A^ transgenes inserted at identical genomic positions. For these experiments we used the UASp promoter (Rørth, 1998), which expresses at lower levels in somatic tissues than UASt (attempts to generate a transgenic line expressing Kib^S677A^ under the ubiquitin promoter were unsuccessful, presumably because ubiquitous expression of a stabilized form of Kib is lethal). Kib^S677A^-GFP accumulated to much greater levels than wild-type Kib-GFP when expressed in the wing disc pouch using the *nub>Gal4* driver (Figures 3F–3H). Confocal imaging revealed that while Kib-GFP and Kib^S677A^-GFP localized similarly, Kib^S677A^-GFP formed bright foci in the basal sections of the tissue, presumably due to higher protein levels (Figures S3H-S3I’). Consistent with the observed increased protein abundance, expression of Kib^S677A^-GFP under the *nub>Gal4* driver led to significantly smaller adult wings than did wild-type Kib-GFP (Figures 3I and 3J). Collectively, these results indicate that Slimb regulates Kib stability *in vivo*.

### Hippo pathway regulates Kibra abundance via Slimb

To this point, our results identify both the Hippo pathway and Slimb as regulators of Kib abundance, but they do not resolve whether the two mechanisms act in parallel or together. We reasoned that if Slimb regulates Kib levels in parallel to the Hippo pathway, then inactivation of the Hippo pathway in tissue expressing Kib^S677A^ would have an additive effect on Kib levels. Conversely, if the Hippo pathway regulates Kib abundance via Slimb, then Kib^S677A^ should be insensitive to pathway inactivation. We first tested the effect of reducing Hippo pathway activity on ubiquitination of Kib^S677A^. In striking contrast to wild-type Kib, ubiquitination of Kib^S677A^ was not sensitive to depletion of Hpo and Wts (Figure 3E), suggesting that Hippo pathway activity promotes Kib degradation via Slimb-mediated ubiquitination.

To test if the Hippo pathway promotes Kib degradation via Slimb *in vivo*, we induced *Mer* mutant clones in wing imaginal discs expressing either wild-type Kib-GFP or Kib^S677A^-GFP under the *nub>Gal4* driver. Similar to endogenous Kib (Figure 1B) or Kib expressed by the ubiquitin promoter (Figure 2B), UASp-Kib-GFP was dramatically upregulated apically and basally in *Mer* clones relative to control cells (Figures 4A–4C’, quantified in 4G). In contrast, Kib^S677A^-GFP appeared only mildly apically stabilized in *Mer* clones (4D-4E’), with no detectible difference in basal Kib^S677A^-GFP levels between the clone and control cells (Figures 4F and 4F’, quantified in 4H), suggesting that the Hippo pathway regulates Kib levels via the degron motif.

**Figure 4:**
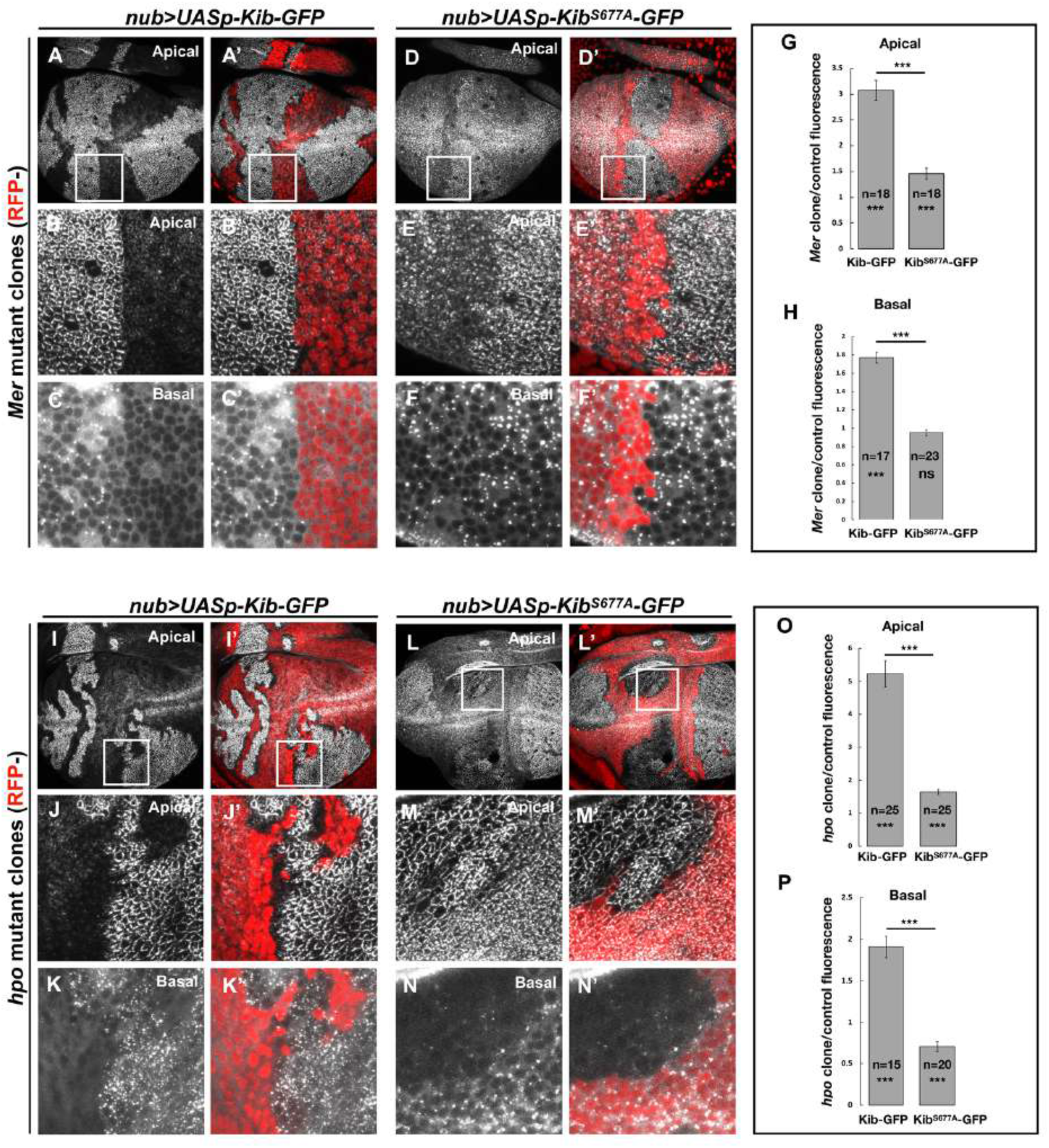
The Hippo pathway regulates Kibra abundance via a putative degron motif. A-F’) *Mer* somatic mosaic clones in wing discs expressing either UASp-Kib-GFP (A – C’) or UASp-Kib^S677A^-GFP (D-F’) with the *nub>Gal4* driver. Note that wild-type Kib-GFP is significantly elevated in *Mer* clones both apically and basally, while Kib^S677A^-GFP is only slightly stabilized apically and is not affected basally. G-H) Quantification of clone/control ratio of apical (G) and basal (H) Kib-GFP fluorescence. All quantification is represented as the mean ± SEM; asterisks above the plots show p-values between the transgenes; asterisks inside each bar show p-values for each transgene with respect to 1; n = number of clones; \*\*\**p* ≤ 0.001, ns = not significant. I-N’) *hpo* somatic mosaic clones in wing discs expressing either UASp-Kib-GFP (I-K’) or UASp-Kib^S677A^-GFP (L-N’) with the *nub>Gal4* driver. Note that whereas wild-type Kib-GFP levels are significantly elevated in *hpo* clones both apically and basally, Kib^S677A^-GFP is stabilized apically but depleted basally in *hpo* clones. O-P) Quantification of clone/control ratio of apical (O) and basal (P) Kib-GFP fluorescence.

The slight apical stabilization of Kib^S677A^-GFP in *Mer* clones could be caused by two possibilities that are not mutually exclusive: 1) Slimb could still weakly bind Kib^S677A^-GFP and promote its degradation, albeit with reduced efficiency, and 2) Hippo pathway activity could inhibit Kib accumulation at the junctional cortex, in which case Hippo pathway inactivation would result in greater cortical Kib accumulation at the expense of the cytoplasmic pool. In support of the first possibility, Kib^S677A^-GFP weakly associated with Slimb (Figure 3D) and was still slightly ubiquitinated in S2 cells (Figure 3E). To ask whether the mild junctional accumulation of Kib^S677A^-GFP in *Mer* clones could also be caused by cortical recruitment, we examined Kib in tissues lacking Hpo, which resulted in stronger junctional accumulation of Ubi>Kib-GFP than loss of Mer (Figure S1F). Strikingly, whereas wild-type Kib-GFP increased both junctionally and basally in *hpo* clones (Figures 4I–4K’, quantified in 4O), Kib^S677A^-GFP increased junctionally but decreased basally in *hpo* clones (Figures 4L–4N’, quantified in 4P). These results suggest that the stabilization of Kib^S677A^-GFP observed upon Hippo pathway inactivation is, at least in part, due to the recruitment of Kib to the junctional cortex where it might be stabilized in a protein complex.

### The Hippo pathway promotes Kibra degradation in a highly compartmentalized manner and independently of pathway activation by Expanded

Previous work showed that Ex interacts with Kib in S2 cells and suggested that Kib and Ex function in a complex to regulate the Hippo pathway (Yu et al., 2010; Genevet et al., 2010). In contrast, *in vivo* studies suggest that Kib functions in parallel to Ex and its partner Crb to regulate activity of the downstream kinase cascade (Baumgartner et al., 2010; Yu et al., 2010; Su et al., 2017). Given these observations, we wondered whether loss of Ex would result in elevated Ubi>Kib-GFP abundance similar to the loss of Mer, Sav, Hpo, or Wts. To our surprise, depletion of Ex and Crb, either individually or together, had no detectable effect on Ubi>Kib-GFP levels (Figures 5A–5C’), suggesting that Kib degradation is promoted specifically via Kib-mediated Hippo pathway activity. Moreover, reducing Hippo pathway activity by other means, such as by overexpressing Dachs or depleting Fat, Ds, Echinoid, or Hpo activator Tao-1 (Boggiano et al., 2011), similarly had no effect on Ubi>Kib-GFP levels (Figures S4A-S4F). On the other hand, knockdown of Mats or Kib’s binding partner, Pez (Poernbacher et al., 2012), increased Ubi>Kib-GFP levels (Figures S4G-S4H). These results suggest that the upstream regulation of the Hippo pathway is highly compartmentalized and that the activity of the core kinases is tightly associated with its distinct upstream regulators in a manner that allows little crosstalk between them.

**Figure 5.**
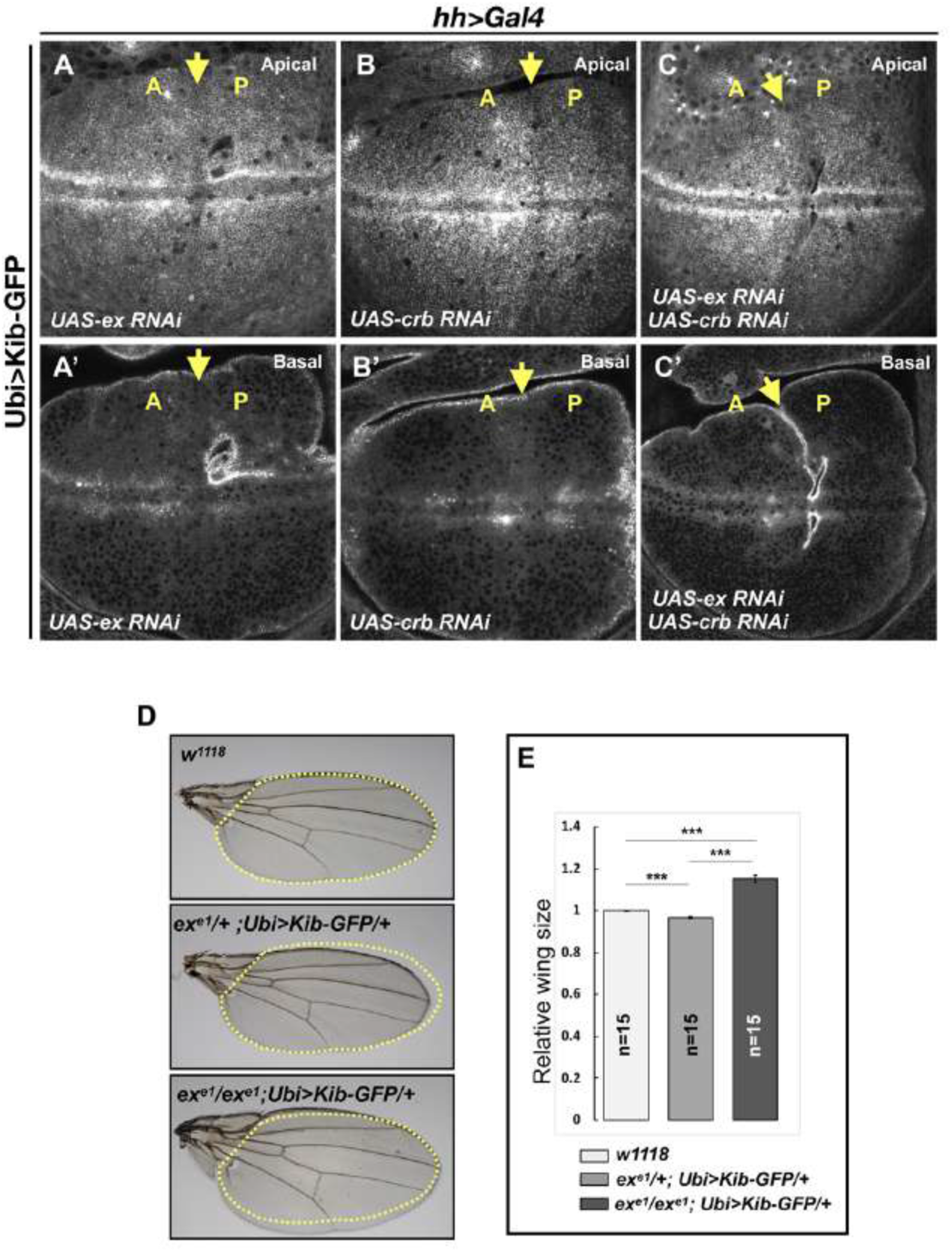
Kibra abundance is regulated independent of Expanded. A-C’) Depletion of Ex (A & A’), Crb (B & B’), or both Ex and Crb (C & C’) in the posterior wing imaginal disc does not affect Ubi>Kib-GFP abundance. Yellow arrows indicate the anterior-posterior (A-P) boundary. D-E) Adult wings of *w^1118^, ex^e1^/+; Ubi>Kib-GFP/+*, or *ex^e1^/ex^e1^; Ubi>Kib-GFP/+* flies. Quantification of wing sizes in (E) is represented as the mean ± SEM; n = number of wings; \*\*\**p* ≤ 0.001.

This parallel behavior of Hippo pathway regulation prompted us to ask whether increasing the activity of one upstream branch of the pathway can substitute for the loss of another. To test this idea, we asked if the Ubi>Kib-GFP transgene, which causes mild undergrowth in a wild-type background (Figure S1E), can suppress the lethality of *ex^e1^*, a null allele. Ubi>Kib-GFP strongly suppressed *ex^e1^* lethality, producing viable and fertile adult flies at expected frequencies (Figure S4J) that completely lacked Ex protein (Figures S4I-S4j’). Homozygous *ex^e1^* flies had significantly larger wings than heterozygotes (Figures 5F-G), but otherwise were phenotypically normal. Together, these results establish that Kib and Ex signal in parallel to regulate at least some aspects of pathway activity.

### The WW domains of Kibra are essential for its degradation via the Hippo pathway and Slimb

Our discovery that Kib degradation is tightly compartmentalized suggests that complex formation between Kib and other Hippo pathway components might be an important step both for pathway activation and Kib degradation. Indeed, Kib interacts with Sav, Mer, Hpo (via Sav) and Wts in S2 cells (Baumgartner et al., 2010; Genevet et al., 2010; Yu et al., 2010) and can recruit these components to the apical cortex *in vivo* (Su et al., 2017). Therefore, we hypothesized that Kib domains that mediate complex formation might also be essential for its degradation.

To test this hypothesis, we performed a structure/function analysis to map the region in Kib essential for its degradation by the Hippo pathway. Kib is a multivalent adaptor protein that contains at least seven potential functional regions: two N-terminal WW domains (WW1 and WW2), a C2-like domain, a putative aPKC-binding domain, and three coiled-coil regions (CC1, CC2, and CC3; Figure 6A). We generated transgenic fly lines expressing different truncations of Kib-GFP under control of the ubiquitin promoter. Two transgenes, one expressing Kib lacking the C2-like domain and another encoding the first 483 amino acids (aa) of Kibra, produced sterile transformants and could not be maintained as stable lines. The rest of the transgenes produced viable and fertile flies.

**Figure 6.**
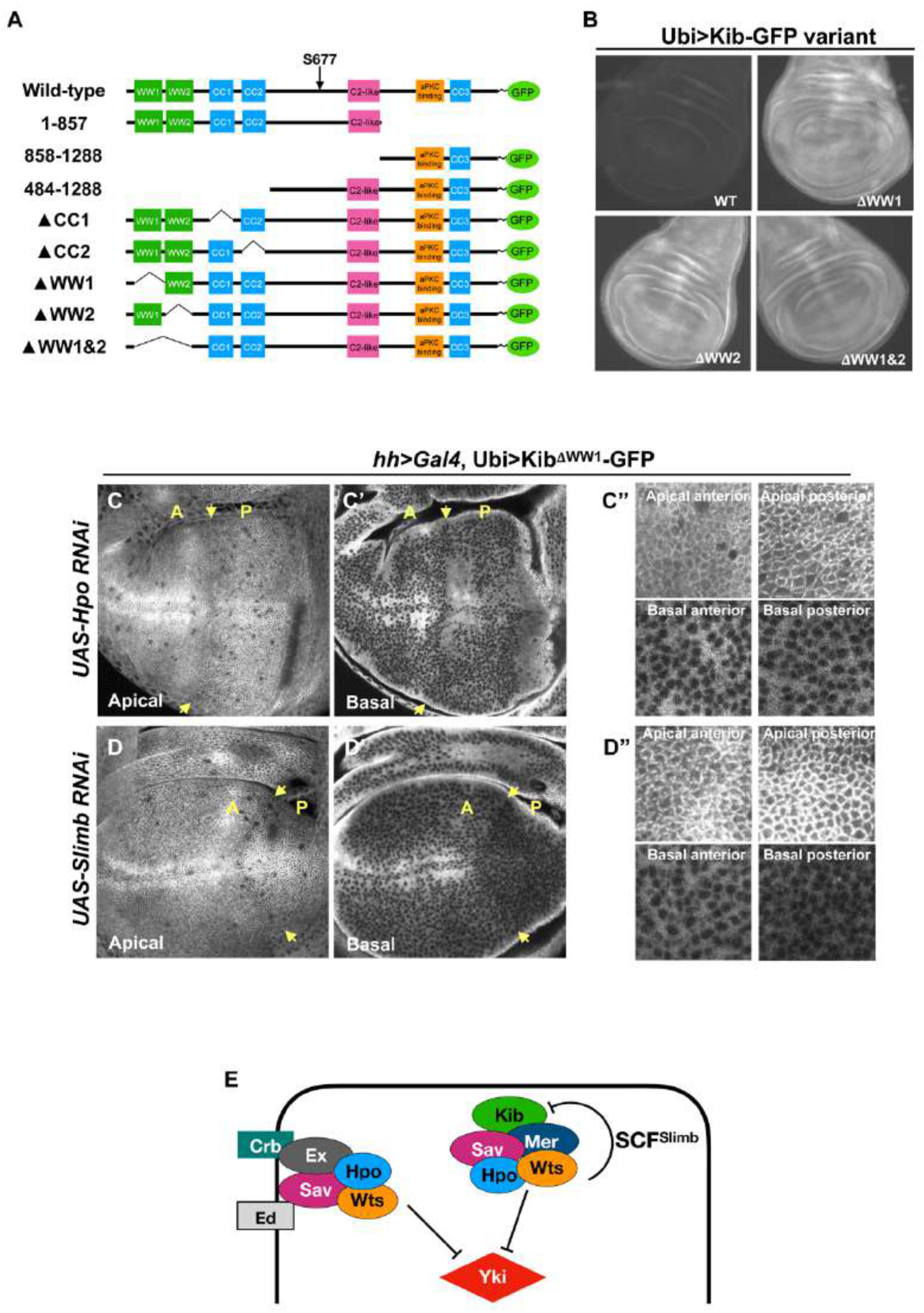
The WW domains of Kib are required for Hippo pathway- and Slimb-mediated degradation. A) Diagram of Kib truncations generated for this study. B) Widefield fluorescence images of wing imaginal discs expressing wild-type and WW-domain truncations of Kib-GFP expressed under the ubiquitin promoter. All images were taken with identical settings. C-C”) Depletion of Hpo does not affect expression of Ubi>Kib^DWW1^-GFP. Note that Hpo depletion leads to apical stabilization and basal depletion of Ubi>Kib^DWW1^-GFP(C”). D-D”) Depletion of Slimb does not affect expression of Ubi>Kib^DWW1^-GFP. Note that similar to Hpo depletion, loss of Slimb leads to slight apical stabilization and basal depletion of Ubi>Kib^DWW1^-GFP (D”). Yellow arrows in indicate A-P boundary of the wing discs. E) A model of compartmentalized Kib degradation by the Hippo pathway and Slimb.

A Kib truncation lacking the C-terminal third of the coding sequence (Ubi>Kib^1-857^-GFP) but retaining the degron motif was strongly upregulated upon Hpo depletion, similar to wild-type Kib (Figures S5A-S5B’). Flies expressing Ubi>Kib^1-857^-GFP had smaller wings than those expressing wild-type Ubi>Kib-GFP (Figure S5J), suggesting that deletion of the C-terminal region enhances Kib activity. In contrast, a Kib truncation lacking the first 483 aa (Ubi>Kib^484-1288^-GFP) was insensitive to Hpo depletion even though it retained the Slimb degron motif (Figures S5D-S5D’), suggesting that the degron alone is not sufficient for pathway-mediated degradation of Kib. Interestingly, Ubi>Kib^484-1288^-GFP was much less potent at suppressing wing growth compared to wild-type Ubi>Kib-GFP (Figure S5J), indicating that the first 483 amino acids of Kib are also essential for Hippo pathway activation.

The first 483 amino acids of Kib contain two WW domains, as well as CC1 and CC2 regions (Figure 6A). Deletion of either CC1 or CC2 did not prevent Kib upregulation upon Hpo depletion, indicating that these regions do not mediate pathway-dependent Kib degradation (Figure S5E-S5E’). However, Kib variants lacking the WW domains, either individually (Ubi>Kib^ΔWW1^-GFP and Ubi>Kib^ΔWW2^-GFP) or together (Ubi>Kib^ΔWW1&2^-GFP), expressed at markedly higher levels than wild-type Kib (Figure 6B). Additionally, these proteins accumulated at the junctional cortex and appeared to be depleted basally but were not upregulated when Hpo was depleted (Figures 6C–6C”, Figures S5G-S5I’). Thus, the WW domains of Kib are necessary for its degradation via the Hippo pathway. Importantly, while Kib lacking the WW domains interacted with Slimb normally in S2 cells, depletion of Slimb had no effect on Ubi>Kib^ΔWW1^-GFP levels (Figures 6D–6D”), again suggesting that association between Kib and Slimb alone is not sufficient for Kib degradation.

Further characterization of the WW domain truncations revealed differences in subcellular localization and effects on growth. Ubi>Kib^ΔWW1&2^-GFP often had an extremely punctate appearance in imaginal tissues (Figures S5N and S5N’). Adult flies expressing Ubi>Kib^ΔWW1&2^-GFP were homozygous viable and had wings almost the size of *w^1118^* controls (Figure S5J) despite the fact that it expressed at higher levels (Figure 6B). Ubi>Kib^ΔWW2^-GFP also had a punctate appearance, but adults expressing Ubi>Kib^ΔWW2^-GFP had significantly smaller wings than flies expressing wild-type Kib-GFP (Figures S5L, S5L’, and S5J). Deletion of WW1, Ubi>Kib^ΔWW1^-GFP, resulted in a protein that localized at the apical cortex but did not form puncta (Figures S5K and S5K’). Flies expressing Ubi>Kib^ΔWW1^-GFP had wings equal in size to *w^1118^* control (Figure S5J). Taken together, these results indicate that while both WW domains are required for pathway mediated Kib turnover, only the WW1 domain of Kib is necessary for Hippo pathway activation.

We reasoned that if complex formation between Kib and other Hippo pathway components is necessary for Kib degradation, then Slimb might also be a part of this complex. Consistent with this prediction, Slimb forms a complex with Mer, Hpo, and Wts in S2 cells (Figures S6D-S6F). We then asked whether the role of the WW domains in Kib degradation was to mediate Kib interaction with other Hippo pathway components. A previous study found that deletion of both WW domains enhanced Kib interaction with Mer in S2 cells (Baumgartner et al., 2010), a result we confirmed (Figure S6B). Kib interacts with Wts in flies (Genevet et al., 2010; Yu et al., 2010), and mammalian Kibra interacts with Lats2 (a mammalian homolog of Wts) via the WW domains (Xiao et al., 2011). We found that the interaction of Kib^ΔWW1&2^ with Wts was significantly weakened but not completely abolished (Figure S6C). Additionally, it was previously reported that Pez interacts with Kib via the WW domains in S2 cells (Poernbacher et al., 2012), consistent with our *in vivo* observation that loss of Pez leads to higher Kib levels. Collectively, these results suggest that pathway-mediated Kib degradation requires the WW domains of Kib, possibly because these domains mediate Kib interaction with multiple pathway components.

### Kibra degradation is patterned across the wing disc epithelium

We next sought to address the potential significance of Kib degradation by the Hippo pathway and Slimb in regulating Hippo pathway activity during development. Observation of wing discs expressing either UASp-Kib-GFP or UASp-Kib^S677A^-GFP revealed strikingly different patterns of Kib distribution. Wild-type Kib-GFP appeared patchy and more concentrated at the center of the tissue with a marked decrease in fluorescence at the periphery of the wing pouch (Figures 7A and 7A’). In contrast, Kib^S677A^-GFP fluorescence was distributed more uniformly throughout the wing pouch, suggesting that degradation of wild-type Kib is relatively greater at the periphery of the pouch compared to its center (Figures 7B and 7B’).

**Figure 7.**
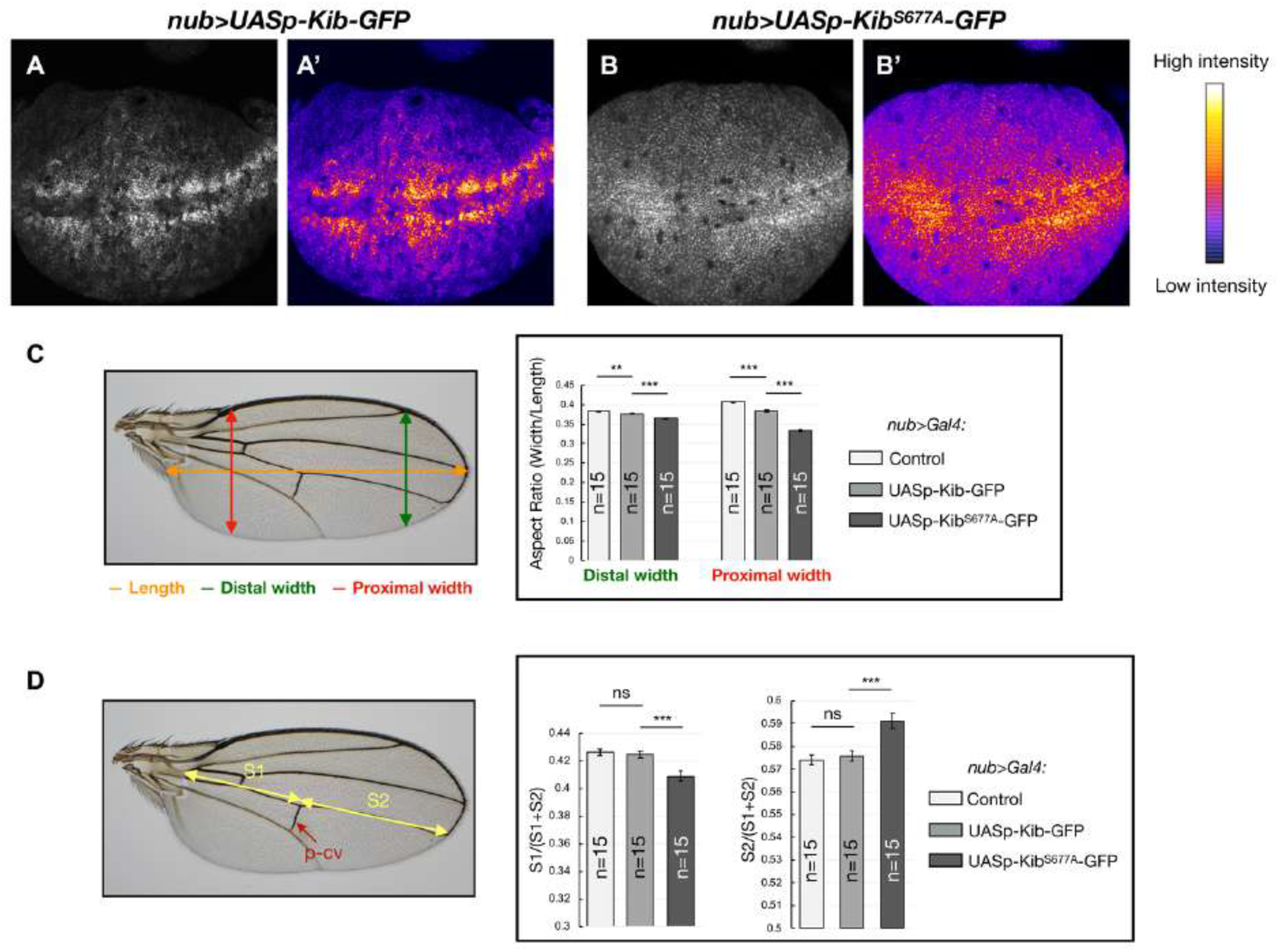
Kib degradation is patterned throughout the wing pouch. A-B’) Grayscale images of the wing pouch, which produces the adult wing blade, expressing UASp-Kib-GFP (A) or UASp-Kib^S677A^-GFP (B) under the *nub>Gal4* driver. Corresponding heatmap intensity images are shown in A’ and B’. Note that Kib^S677A^-GFP displays a more uniform distribution across the pouch than wild-type Kib-GFP. See also Figure S7A. C) Quantification of aspect ratios of adult wings expressing *nub>Gal4* alone or with UASp-Kib-GFP and UASp-Kib^S677A^-GFP. The color-coded segments in the wing image represent the wing length (orange), distal width (green), and proximal width (red). D) Quantification of the length of proximal (S1) or distal (S2) wing region with respect to total wing length in wings expressing *nub>Gal4* alone or with UASp-Kib-GFP and UASp-Kib^S677A^-GFP; p-cv = posterior cross vein. Quantification is represented as the mean ± SEM; n = number of wings; \*\*\**p* ≤ 0.001, \*\**p* ≤ 0.01, ns = not significant.

To ask whether the observed difference in protein distribution affected tissue growth, we analyzed adult wings of w^1118^ flies expressing either UASp-Kib-GFP or UASp-Kib^S677A^-GFP under the *nub>Gal4* driver. The peripheral regions of the wing pouch correspond to the proximal regions of the adult wing. To ask whether growth was disproportionately inhibited in the proximal region of the wing, we first measured the wing aspect ratios comparing the width of the proximal or distal wing regions to the overall proximal-distal length. Strikingly, while the relative decrease in width distally was mild in *nub>*UASp-Kib-GFP or *nub>*UASp-Kib^S677A^-GFP wings compared to control wings, the proximal width decreased dramatically in wings expressing Kib^S677A^, indicating that expression of Kib^S677A^ inhibited growth disproportionately more in the proximal region (Figure 7C). Similarly, when the wing length was measured in the proximal-distal (P-D) axis, using L4 vein as an estimate of total length and the posterior crossvein (p-cv) as the approximated midpoint, we found that wing growth was more severely inhibited proximally than distally (Figure 7D). Collectively, these results suggest that Kib degradation occurs in a patterned manner in the wing imaginal epithelium and could serve to pattern tissue growth.

## Discussion

In this study, we show that the Hippo pathway negatively regulates Kib levels via a previously unrecognized post-translational feedback loop. Several key results indicate that this feedback is independent of Yki transcriptional activity. First, loss of Mer leads to a dramatic increase in Kib levels without a detectable increase in Yki transcriptional activity. Second, removing Sd, which blocks Yki-mediated transcription, does not suppress the elevated Kib levels upon Hippo pathway inactivation. Third, the abundance of Kib-GFP expressed under a Yki-insensitive promoter (Ubi>Kib-GFP) still increases upon Hippo pathway inactivation. Additionally, we show that Hippo pathway activity promotes Kib phosphorylation and ubiquitination and identify a ubiquitin ligase, SCF^Slimb^, that mediates Kib ubiquitination in a pathway-dependent manner.

Multiple upstream components regulate the core Hippo kinase cassette, but their organization and the degree of crosstalk between them has not been well elucidated. A striking aspect of our findings is the extent to which Kib degradation by the Hippo pathway is insulated from the activity of other upstream pathway regulators. Previous studies have shown that Crb, Ex, and Ed function together at the junctional cortex (Sun et al., 2015) and in parallel to Kib at the medial cortex (Su et al., 2017). Ft can influence Ex stability at the junctions (Silva et al., 2006; Wang et al., 2019), suggesting linkage between these Hippo signaling branches, and Tao-1 functions downstream of Ex (Chung et al., 2016). Although Ex can form a complex with Kib in S2 cells (Genevet et al., 2010; Yu et al., 2010) and Kib junctional localization is dependent on Crb (Su et al., 2017), our results show that depletion of Crb, Ex, Ft, Ed, or Tao-1 does not affect Ubi>Kib-GFP level. Combined with our results that the more downstream components Hpo and Wts promote Kib phosphorylation, ubiquitination and degradation, our data strongly suggest that Kib degradation is mediated via Kib-promoted Hippo pathway complex formation. Such tightly compartmentalized activity of the Hippo pathway is also suggested by the punctate appearance of all of these proteins at the cell cortex (Su et al., 2017, Supplemental Figure S1D). While our data are consistent with a model in which Hpo or Wts directly phosphorylates Kib, we do not have direct evidence for this and it is possible that other components are involved. Compartmentalized, parallel regulation of pathway activity could clearly have functional implications for the control of tissue growth but at the moment is poorly understood.

A remaining question our work defines relates to the functional significance of this mechanism to regulate Kib abundance in developing tissues. Signal activation is often tightly coupled to downregulation, for example by endocytosis and degradation of a cell surface receptor, in a manner that ensures that pathway activation can be transient and tightly regulated. Our structure/function analysis revealed a very strong correlation between Kib degradation by the Hippo pathway and the ability of Kib to promote pathway activation, suggesting that activation/inactivation in Hippo signaling is also tightly coupled. We currently know little about how dynamically Hippo pathway output is regulated in developing tissues, largely because there are no available singlecell resolution reporters for pathway activity, but our findings suggest that pathway output can be tightly and dynamically regulated within cells of growing tissues.

Our results also suggest that Kib-mediated Hippo signaling is patterned across the wing imaginal epithelium by regulated Kib abundance. Our observations indicate that whereas wild-type Kib is expressed in a patchy pattern and is more concentrated at the tissue center than the periphery, Kib^S677A^ is distributed more uniformly throughout the tissue. Since both Kib transgenes were identically expressed (from the same genomic location and using the same Gal4 driver), these differences are likely produced via regulation of Kib stability. Previous studies have described marked differences in tension, cell size, and cell shape between the periphery and the center of the wing imaginal tissue (LeGoff et al., 2013; Mao et al., 2013), suggesting that these factors could modulate Kib levels in a region-specific manner. Interestingly, junctional tension is known to regulate the Hippo pathway both in *Drosophila* and mammalian cells via inhibition of Wts (Lats1/2 in mammals) (Rauskolb et al., 2014; Ibar et al., 2018), though it remains unknown whether junctional tension affects upstream Hippo pathway regulators such as Kib. Additionally, higher tension at the wing pouch periphery was proposed to drive tissue growth, possibly by inhibiting the Hippo pathway, as a compensatory mechanism to the lower morphogen levels (Aegerter-Wilmsen et al., 2007; Hariharan, 2015). The results from this study may lead to the elucidation of how external cues, such as mechanical tension, modulate Kib-mediated Hippo signaling and pattern tissue growth.

Another question our study raises is the functional significance of having both transcriptional and post-translational negative feedback mechanisms that regulate Kib levels. Feedback regulation is a common feature in cell signaling, and transcriptional negative feedback can serve to limit the output of a signaling pathway over time (Perrimon and McMahon, 1999). In the case of the Hippo pathway, transcriptional feedback mediated by Yki is not specific to Kib, as Ex and Mer expression is also promoted by Yki activity. Moreover, loss of any upstream Hippo pathway regulator, including Ex, Crb, Ft, or Ed would presumably affect Kib levels via the transcriptional feedback. In contrast, the post-translational feedback identified in this study would silence Kib function in a more rapid and specific manner. The role of the post-translational feedback could be to enhance the robustness of Kib-mediated signaling (Stelling et al., 2004), possibly by preventing drastic fluctuations in Kib levels, to ensure optimally scaled and patterned tissue growth. On a broader level, our identification of Kib-specific feedback highlights the importance of understanding why there are multiple upstream inputs regulating the Hippo pathway and how they function during development.

## Methods

### Fly genetics

For expression of UAS transgenes, the following drivers were used: *hh>Gal4, en>Gal4, ap>Gal4, nub>Gal4.*

To generate mutant clones, the following crosses were performed:

Kib::GFP in *ex* or *Mer* mutant clones:

*y w hsFlp; Ubi-RFP 40A FRT X ex^e1^ 40A FRT/CyO, dfdYFP; Kib::GFP/TM6, Tb Mer^4^ 19A FRT/FM7, actGFP; MKRS/TM3, Ser, actGFP X hsFLP, w^1118^, Ubi-RFP-nls 19AFRT; Kib::GFP/TM3, Ser, actGFP*

Yki-YFP in *ex* or *Mer* mutant clones:

*y w hsFlp; Ubi-RFP 40A FRT X ex^e1^ Yki-YFP yki^B5^/CyO, dfdYFP Mer^4^ 19A FRT/+; Yki-YFP/CyO, dfdYFP X hsFLP, w^1118^, Ubi-RFP-nls 19AFRT;MKRS/TM3, Ser, actGFP*

*ban3-GFP* in *ex* or *Mer* mutant clones:

*y w hsFlp; Ubi-RFP 40A FRT X ex^e1^ 40A FRT/CyO, dfdYFP; ban3-GFP/TM6, Tb Mer^4^ 19A FRT/FM7, actGFP; MKRS/TM3, Ser, actGFP X hsFLP, w^1118^, Ubi-RFP-nls 19AFRT; ban3-GFP/TM3, Ser, actGFP*

Kib::GFP in *sd Mer* double mutant clones:

*sd^47^ Mer^4^ 19A FRT/FM7, dfdYFP; Sco/CyO, dfdYFP X hsFLP, w^1118^, Ubi-RFP-nls 19AFRT; Kib::GFP/TM3, Ser, actGFP*

Ubi>Kib-GFP in single *sd* or *hpo* mutant clones or in *sd hpo* double mutant clones:

*sd^47^ 19A FRT/FM7, dfdYFP; FRT 42D hpo^BF33^/CyO, dfdYFP X ey>Flp Ubi-GFP 19A FRT; FRT 42D Ubi-RFP/CyO, dfdYFP; Ubi>Kib-GFP/+*

UASp-Kib-GFP or UASp-Kib^S677A^-GFP in *Mer* or *hpo* mutant clones:

*Mer^4^ 19A FRT/+; nub>Gal4/CyO, dfdYFP X hsFLP, w^1118^, Ubi-RFP-nls 19AFRT; UASp-Kib-GFP/TM3, Ser, actGFP*

*Mer^4^ 19A FRT/+; nub>Gal4/CyO, dfdYFP X hsFLP, w^1118^, Ubi-RFP-nls 19AFRT; UASp-Kib^S677A^-GFP/TM3, Ser, actGFP*

*nub>Gal4 FRT 42D hpo^BF33^/CyO, dfdYFP X y w hsFLP; FRT 42D Ubi-RFP/CyO, dfdYFP; UASp-Kib-GFP/+*

*nub>Gal4 FRT 42D hpo^BF33^/CyO, dfdYFP X y w hsFLP; FRT 42D Ubi-RFP/CyO, dfdYFP; UASp-Kib^S677A^ -GFP/+*

### Expression constructs and generation of *Drosophila* transgenic lines

To generate Ubi>Kib-GFP, Kib was fused to GFP-FLAG with a linker sequence 5’-TCCGGTACCGGCTCCGGC-3’, and the entire Kib-GFP-FLAG cassette was first cloned into UAStattB backbone (Bischof et al., 2007) to generate UASt-Kib-GFP-FLAG, with unique NotI (immediately 5’ of the Kozak sequence) and KpnI (in the linker region) restriction sites flanking Kib sequence. To make Kib^1-857^, Kib^858-1288^, Kib^484-1288^, the corresponding regions were amplified (see primers table); UAStattB was linearized with NotI and KpnI and the amplified fragments were cloned into linearized backbone via Gibson assembly (Gibson et al., 2009). Fragments lacking CC or WW domains were made using an inverse PCR approach with flanking primers (see primers table) and the amplified linear pieces including the plasmid backbone were circularized via Gibson assembly. Kib-GFP-FLAG cassettes (full-length or truncations) were amplified using flanking primers (see table) and cloned via Gibson assembly into *p63E-ubiquitin* backbone (Munjal et al., 2015) linearized with NotI and XbaI. The transgenes were inserted at the 86Fb (full-length Kib) or VK37 (full-length and truncated Kib) docking site via phiC31-mediated site-specific integration.

pMT-Kib-GFP-FLAG was generated by cloning Kib-GFP-FLAG cassette via Gibson assembly (Gibson to pMT primers) into the pMT backbone (Klueg et al., 2002) linearized by KpnI and EcoRV.

UASp-Kib^S677A^-GFP-FLAG was generated using Q5^®^ Site-Directed Mutagenesis Kit (New England Biolabs, Catalog #E0554S) using primers KibS677A (see primers table). pMT-Kib-GFP-FLAG was used as a template due to smaller size of the plasmid. The mutant Kib^S677A^-GFP-FLAG cassette was excised with NotI and XbaI and ligated into pUASp (Rørth, 1998) to generate UASp-Kib^S677A^-GFP. Both UASp-Kib^S677A^-GFP and UASp-Kib-GFP were inserted at the 86Fb docking site via phiC31-mediated sitespecific integration.

### Immunostaining of imaginal tissues

Wing or eye imaginal discs from wandering late third instar larvae were fixed and stained as previously described (McCartney and Fehon, 1996). Primary antibodies, listed in Key Resources table, were diluted as follows: anti-Ex (1:5000), anti-FLAG (1:20,000), anti-Sd (1:1000). Secondary antibodies (diluted 1:1000) were from Jackson ImmunoResearch Laboratories. Immunostaining samples were imaged using either a Zeiss LSM 800 or LSM 880 confocal microscope and the images were analyzed with *Image J.*

### Live imaging of imaginal tissues

Live imaging of the *Drosophila* imaginal tissues was performed as previously described (Xu et al., 2019). Briefly, freshly dissected wing or eye imaginal discs from third instar larvae were pipetted into a ~40μl droplet of Schneider’s *Drosophila* Medium supplemented with 10% Fetal Bovine Serum and mounted on a glass slide. To support the tissue, spherical glass beads (Cospheric, Product ID: SLGMS-2.5) of ~50μm in diameter were place under the cover slip. The mounted samples were immediately imaged on Zeiss LSM 880 or LSM 800 confocal microscopes. Widefield fluorescence imaging of live wing imaginal discswas done using a Zeiss Axioplan 2ie microscope with an Orca ER camera.

### Co-immunoprecipitation from S2 cells

The following constructs were used in co-IP experiments: *pMT-Kib-GFP-FLAG* (this study), *pMT-Kib^ΔWW1&2^-GFP-FLAG* (this study)*, pAc5.1-Slimb-6x-myc* (from J. Chiu, UC Davis), *pAFW-Mer, pAHW-Mer^1-600^, pMT-FLAG-Hpo*, and *pAC5.1-V5-Wts* (Huang et al., 2005).

Briefly, 3.5 x 10^6^ S2 cells were transfected with total of 500ng of the indicated DNA constructs using DDAB (dimethyldioctadecylammonium bromide, Sigma) (Han, 1996) at 250 μg/ml in six-well plates. Immunoprecipitation (IP) was performed 3 days after transfection. For expression of pMT constructs, 700 μM CuSO4 was added to the wells 24h prior to cell lysis (2 days after transfection). For GFP or Myc IPs, guinea pig anti-GFP (1:1,250) or mouse anti-Myc (1:1,000) antibodies were used. Pierce™ Protein A (Thermo Scientific™) magnetic beads were used to precipitate antibody-bound target proteins. For immunoblotting the following antibody concentrations were used: rabbit anti-GFP (1:5,000), mouse anti-Hpo (1:5,000), mouse anti-a-tubulin (1:2,500), mouse anti-Myc (1:40,000), mouse M2 anti-Flag (1:20,000), mouse anti-V5 (1:2,500), and rabbit anti-HA (1:5,000). Immunoblots were scanned using an Odyssey CLx scanner (LI-COR Biosciences).

Cells were harvested and lysed on ice in buffer containing 25 mM Hepes, 150 mM NaCl, 1 mM EDTA, 0.5 mM EGTA, 0.9 M glycerol, 0.1% Triton X-100, 0.5 mM DTT, and Complete protease inhibitor cocktail (Roche) at 1 tablet/10ml concentration.

For detection of phosphorylated Kib *in vivo*, wing discs from wandering third-instar larvae (200 discs per condition) expressing *nub>Gal4* with Ubi>Kib-GFP alone or together with an indicated RNAi transgene were dissected and immediately flash-frozen in a bath of dry ice and 95% ethanol (stored at −80°C). On the day of IP, the samples were briefly thawed on ice and lysed in buffer described above. PhosSTOP™ (Sigma) phosphatase inhibitor cocktail was added to the lysis buffer to inhibit phosphorylation (1 tablet/10ml of buffer). Kib-GFP was immunoprecipitated with guinea-pig anti-GFP antibody (1:1,250). A control sample was treated with λ-phosphatase. Samples were run on 8% polyacrylamide gel, with 118:1 acrylamide/bisacrylamide (Scheid et al., 1999), to better resolve phosphorylated Kib species.

### Detection of ubiquitinated Kibra in S2 cells

For ubiquitination assays, pMT-HA-Ub (Zhang et al., 2006) was co-transfected where indicated to provide labeled ubiquitin. To inhibit proteasomal degradation, 50μM MG132 (Cayman Chemical) and 50μM calpain inhibitor I (Sigma) was added 4h prior to cell lysis. Cells were lysed in RIPA buffer (150mM NaCl, 1%NP-40, 0.5% Na deoxycholate, 0.1%SDS, 25mM Tris (50mM, pH 7.4) supplemented with 5 mM N-Ethylmaleimide *(NEM)*, and Complete protease inhibitor cocktail (Roche, 1 tablet/10ml of buffer). HA-tagged ubiquitin was purified using Pierce anti-HA Magnetic beads (clone 2-2.2.14).

### Quantification and statistical analysis

*Image J* was used to quantify mean fluorescence intensity in clones vs. control region in Figs. 1C, 1H, 4G, 4H, 4O, and 4P. In all cases, no more than two clones per imaginal disc were used for quantification. To quantify adult wing sizes, wings were mounted in methyl salicylate and photographed with the same settings on a Zeiss Axioplan 2ie microscope using a Canon camera (EOS rebel T2i). Subsequent measurements of wing size and statistical analyses were processed using Image J and MS Excel, respectively. All statistical comparisons between means were performed using Student’s t-test. Statistical significance of results between compared groups was indicated as follows: *** (p ≤ 0.001), ** (p ≤ 0.01), * (p ≤ 0.05), ns (not significant, p > 0.05).

## Author Contributions

S. A. T, T. S, and R. G. F conceived the project. T. S. made the initial observations of the effect of loss of Slimb and Hippo pathway activity on Kib levels and generated plasmids with Kib truncations. A. U. optimized and performed the ubiquitination assays. S. A. T. performed all other experiments and generated all other reagents. S. A. T. and R. G. F. analyzed data and wrote the manuscript. R. G. F. supervised all aspects of the project.

## Acknowledgements

We thank D. Pan, K. Irvine, J. Jiang, J. Chiu, M. Glotzer, T. Lecuit, K. Guss, the Developmental Studies Hybridoma Bank, TRiP at Harvard Medical School (NIH/NIGMS R01-GM084947), the Bloomington stock center, and VDRC stock center for fly stocks and other reagents. We thank S. Buiter for technical help and Y. Wang for help generating the guinea pig anti-GFP antibody. We thank M. Glotzer and S. Horne-Badovinac for helpful comments on the manuscript. S. A. T. was supported by an NIH training grant (T32 GM007183) and an NSF-GRFP. This work was supported by a grant from the National Institutes of Health to R.G.F (NS034783).

