## Supplemental Figures for "Yorkie-independent negative feedback couples Hippo pathway activation with Kibra degradation"

### Supplemental Figure S1

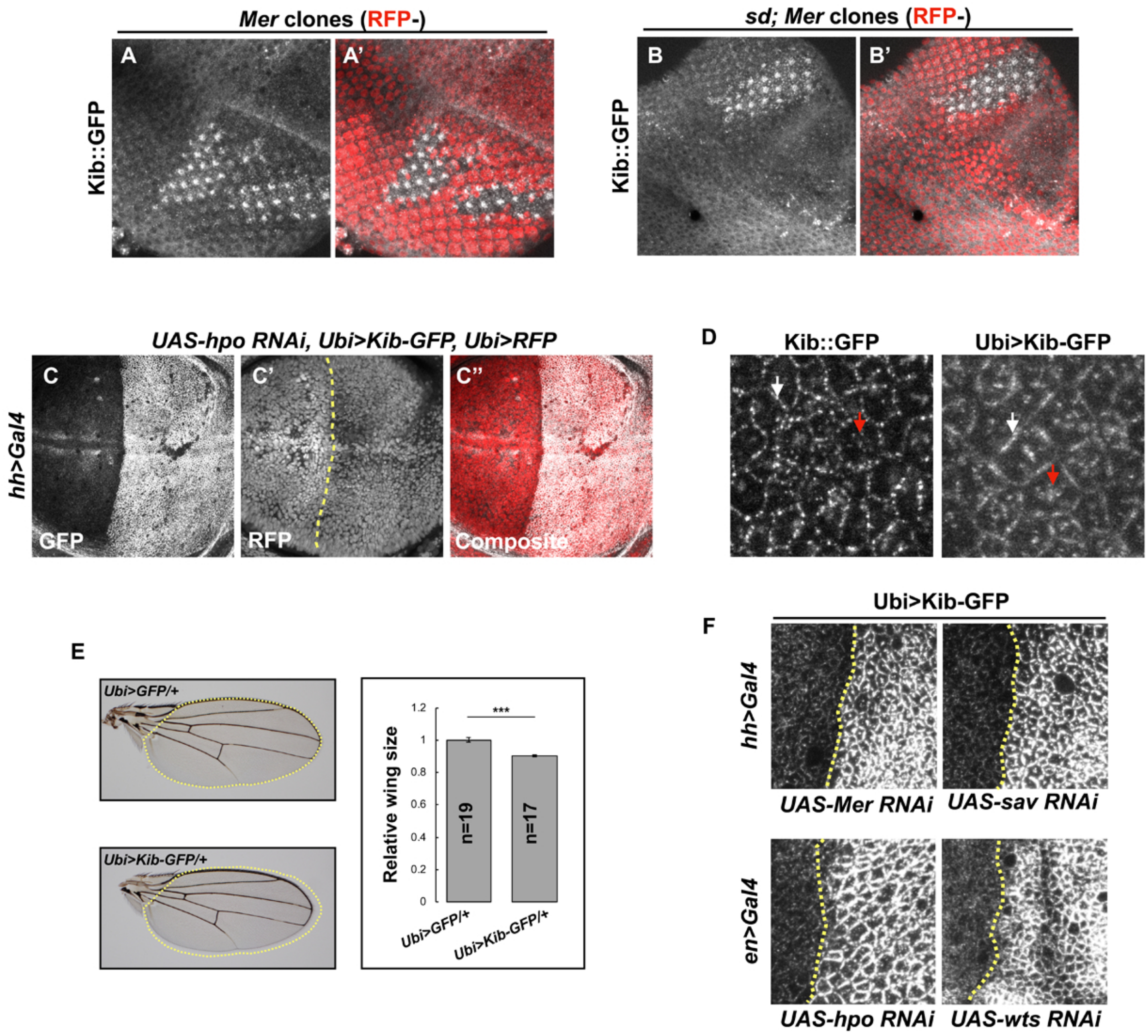

**Figure S1, related to Figure 2.**

**The Hippo pathway regulates Kib levels independent of Yki transcriptional activity.**

A-B') Endogenous Kib:GFP levels are elevated in *Mer* somatic mosaic clones (A & A') and in double *Mer; sd* clones (B & B').

C-C'') Depletion of Hpo in the posterior compartment of the wing does not cause increased Ubi>RFP expression. Yellow dashed line indicates the anterior-posterior (A-P) boundary.

D) Similar to endogenous Kib:GFP (left), Ubi->Kib-GFP (right) accumulates both at the junctional (white arrows) and apical medial cortex (red arrows).

E) Size comparison of adult wings from flies expressing Ubi>GFP or Ubi>Kib-GFP; quantification is shown as the mean ± SEM relative to the control; n = number of wings; ***$p \leq 0.001$.

F) Magnified images from Figure 2 showing that Hippo pathway inactivation results in junctional Kib accumulation. Yellow dashed lines mark the A-P boundaries, with posterior to the right.

### Supplemental Figure S2

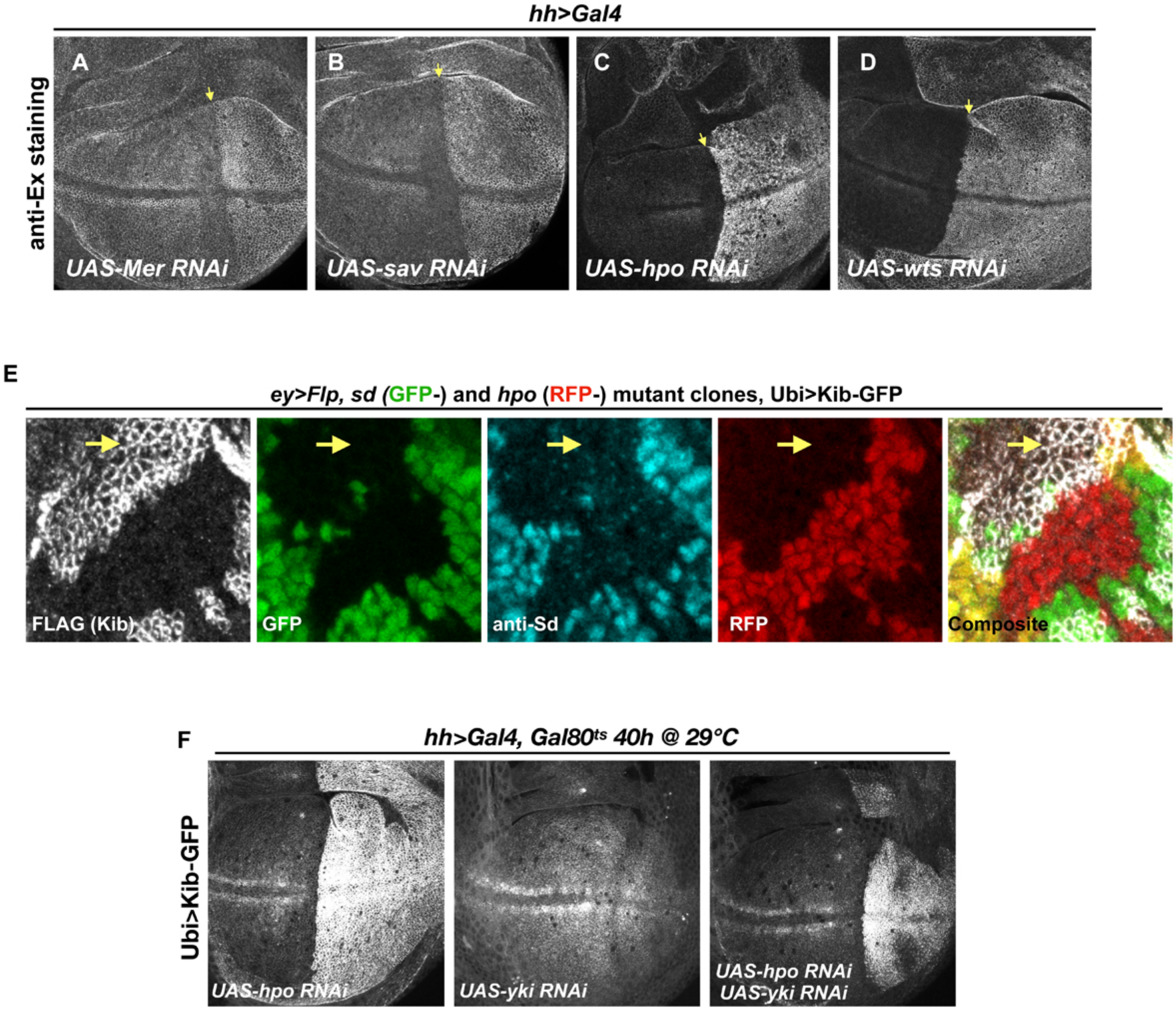

**Figure S2, related to Figure 2.**

**The Hippo pathway regulates Kib levels independent of Ex.**

A-D) Ex levels are elevated upon Hippo pathway inactivation, with a particularly strong increase upon Hpo or Wts depletion (C & D, respectively).

E) Single *sd* (GFP-) or *hpo* (RFP-) somatic mosaic clones or double *sd; hpo* clones (GFP- and RFP-, yellow arrow) induced in the eye imaginal disc using *ey>Flp.* Ubi>Kib-GFP (FLAG staining) is upregulated in *sd; hpo* double mutant clones; loss of *sd* was confirmed by anti-sd staining (cyan).

F) Effect of transient depletion of Hpo (left), Yki (middle), or Hpo and Yki (right) on Ubi>Kib-GFP levels in the posterior compartment of the wing using $Gal80^{ts}$.

### Supplemental Figure S3

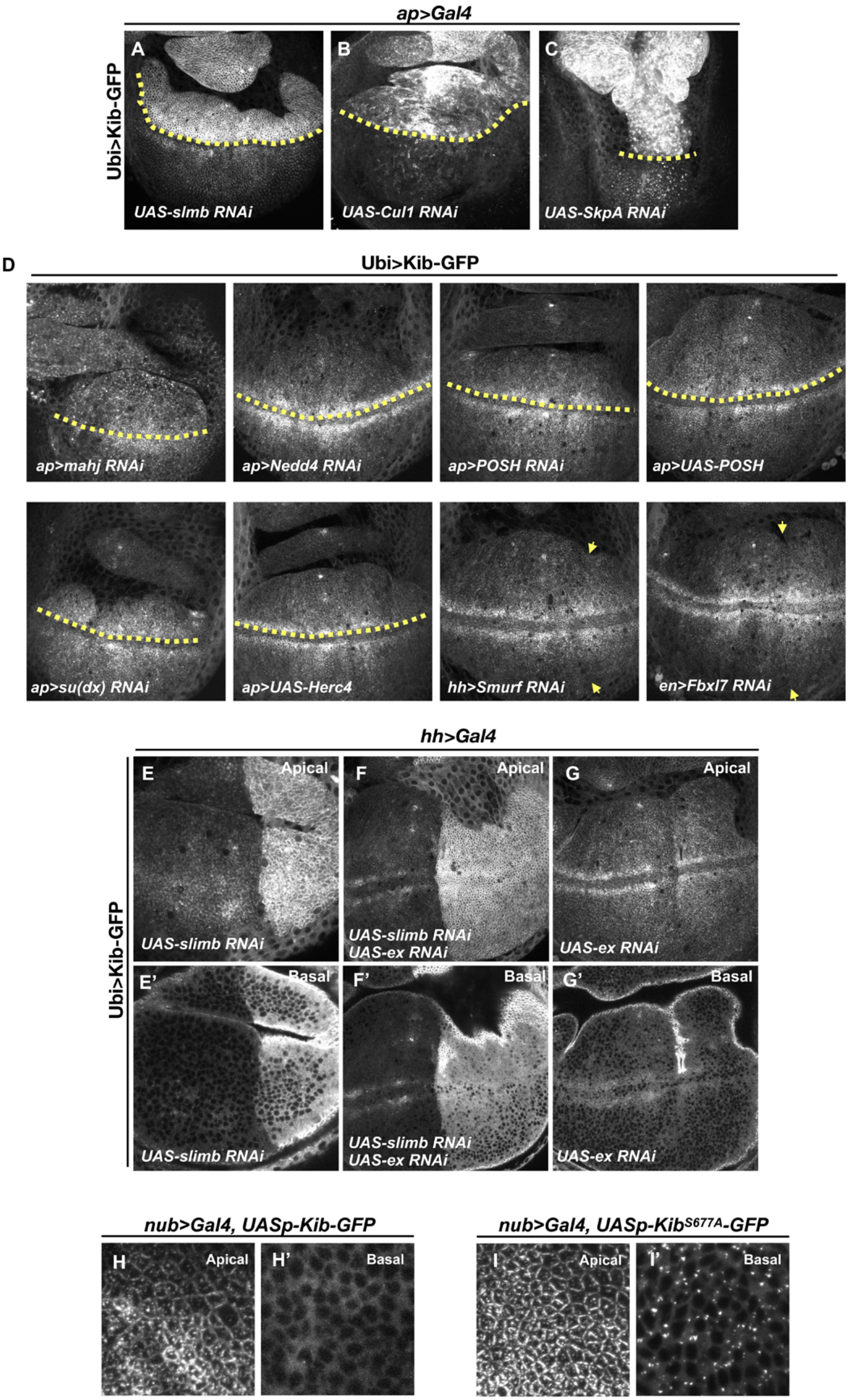

**Figure S3, related to Figure 3.**

**Effect of different E3 ubiquitin ligases involved in the Hippo pathway on Kib levels.**

A-C) Depletion of $SCF^{Slimb}$ E3 ubiquitin ligase components Slimb (A), Cul1 (B), or SkpA (C) in the dorsal compartment of the wing imaginal disc results in increased Ubi>Kib-GFP levels.

D) Depletion or overexpression of other E3 ubiquitin ligases known to regulate Hippo pathway components has no effect on Ubi>Kib-GFP levels.

Yellow dashed line represents the dorsal-ventral boundary, with dorsal side up (for ap>Gal4); yellow arrows indicate the anterior-posterior boundary, with posterior to the right (for hh and en>Gal4).

E-G') Ubi>Kib-GFP levels are elevated upon depletion of Slimb alone or co-depletion of Slimb and Ex, but not when Ex alone is depleted in the posterior compartment of the wing imaginal disc.

H-I') Confocal images of UASp-Kib-GFP or UASp-$Kib^{S677A}$-GFP (using different settings to adjust for difference in abundance) expressed with nub>Gal4. Note that $Kib^{S677A}$-GFP shows similar localization to Kib-GFP apically but $Kib^{S677A}$-GFP forms foci basally.

### Supplemental Figure S4

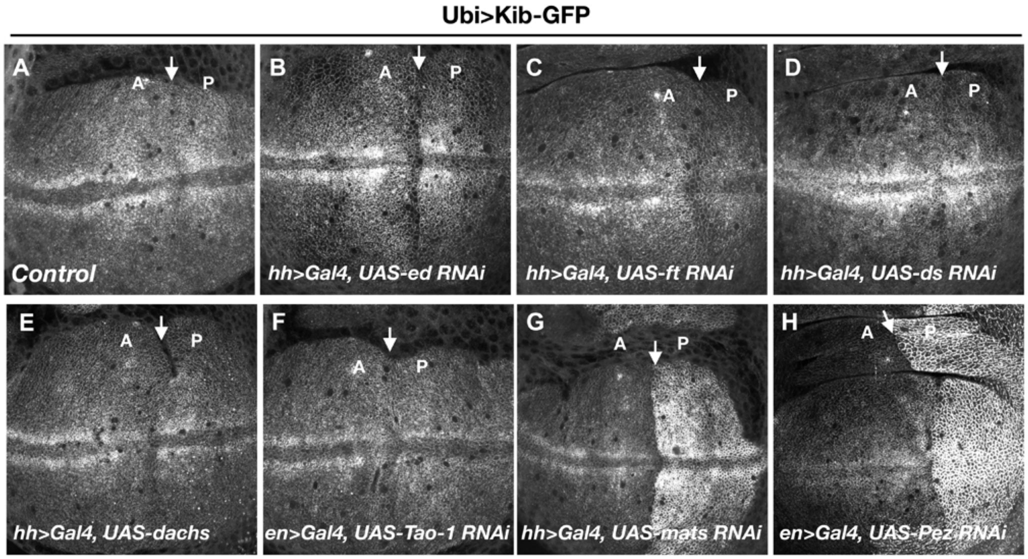


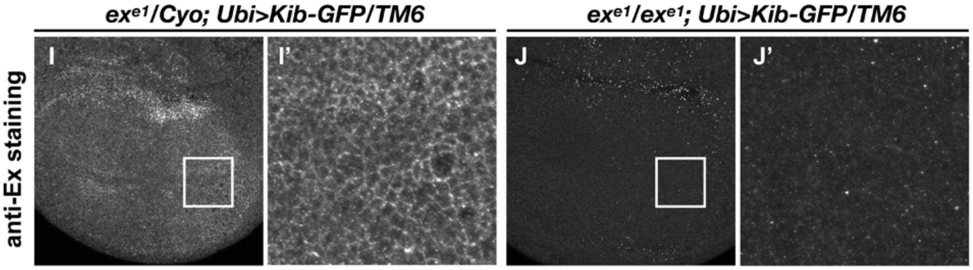


**K**

***$ex^{e1}$/CyO; Ubi>Kib-GFP/TM6  X  $ex^{e1}$/CyO; Ubi>Kib-GFP/TM6***

| $F_1$ progeny | Observed | Expected |
|---|---|---|
| ***$ex^{e1}$/CyO; Ubi>Kib-GFP/TM6*** | 55 | 51 |
| ***$ex^{e1}$/$ex^{e1}$; Ubi>Kib-GFP/TM6*** | 21 | 25 |

$X^2 = 0.95$

$p = 0.31$

**Figure S4, related to Figure 5.**

**The Hippo pathway controls Kib abundance in a tightly compartmentalized manner.**

A-H) Ubi>Kib-GFP levels are regulated by a subset of Hippo pathway components.

I-J') Ectopic Kib suppresses $ex^{e1}$ lethality. Wing discs of $ex^{e1}$ heterozygous larvae (I & I') or $ex^{e1}$ homozygous larvae (J & J') carrying Ubi>Kib-GFP were stained for Ex to confirm the presence of the $ex^{e1}$ allele.

K) Chi-square analysis shows that $ex^{e1}$ homozygotes survive as expected if ectopic Kib completely suppresses *ex* lethality.

### Supplemental Figure S5

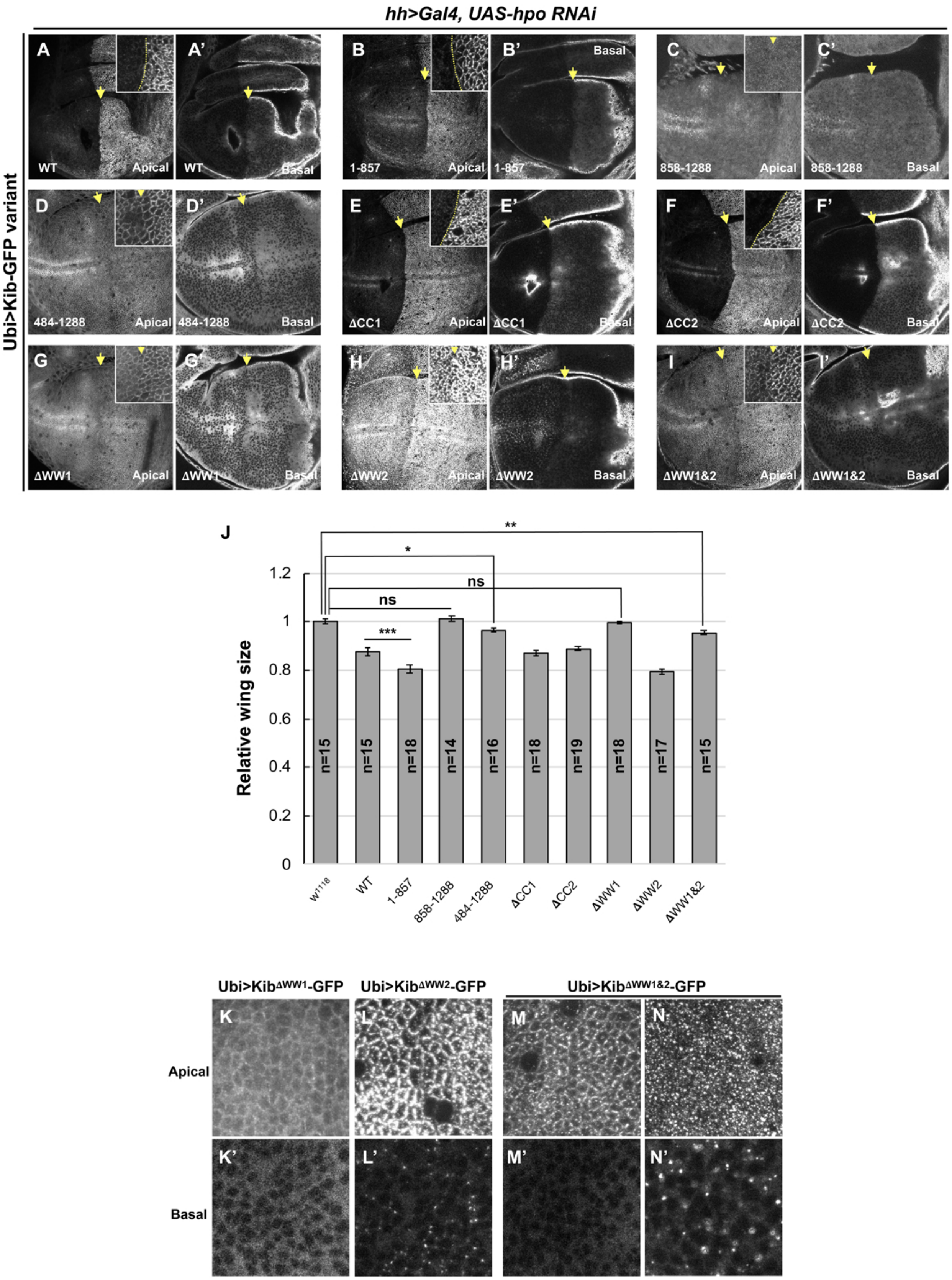

**Figure S5, related to Figure 6.**
**The role of WW domains in Hippo pathway-mediated Kib degradation.**

A-I') Effect of Hpo depletion in the posterior compartment of the wing imaginal disc on different Kib truncations. Deletion of the WW domains, individually (G-H') or together (I & I') stabilizes Kib apically but does not lead to an increase in basal Kib levels. Note that tissue in G is the same as shown in Figure 6C.

J) Size comparison (relative to wild-type) of adult wings from flies ectopically expressing different Ubi>Kib-GFP truncations. Quantification is shown as the mean ± SEM; n = number of wings; ***$p \leq 0.001$, **$p \leq 0.01$, *$p \leq 0.05$, ns = not significant.

K-N') Localization of Kib lacking WW1 (K & K'), WW2 (L & L') or both WW 1&2 (M-N') in wing imaginal disc cells. Note that Ubi>Kib$^{\Delta WW1\&2}$-GFP sometimes localizes normally at the junctions (M), but usually accumulates in bright foci both apically and basally (N and N').

### Supplemental Figure S6

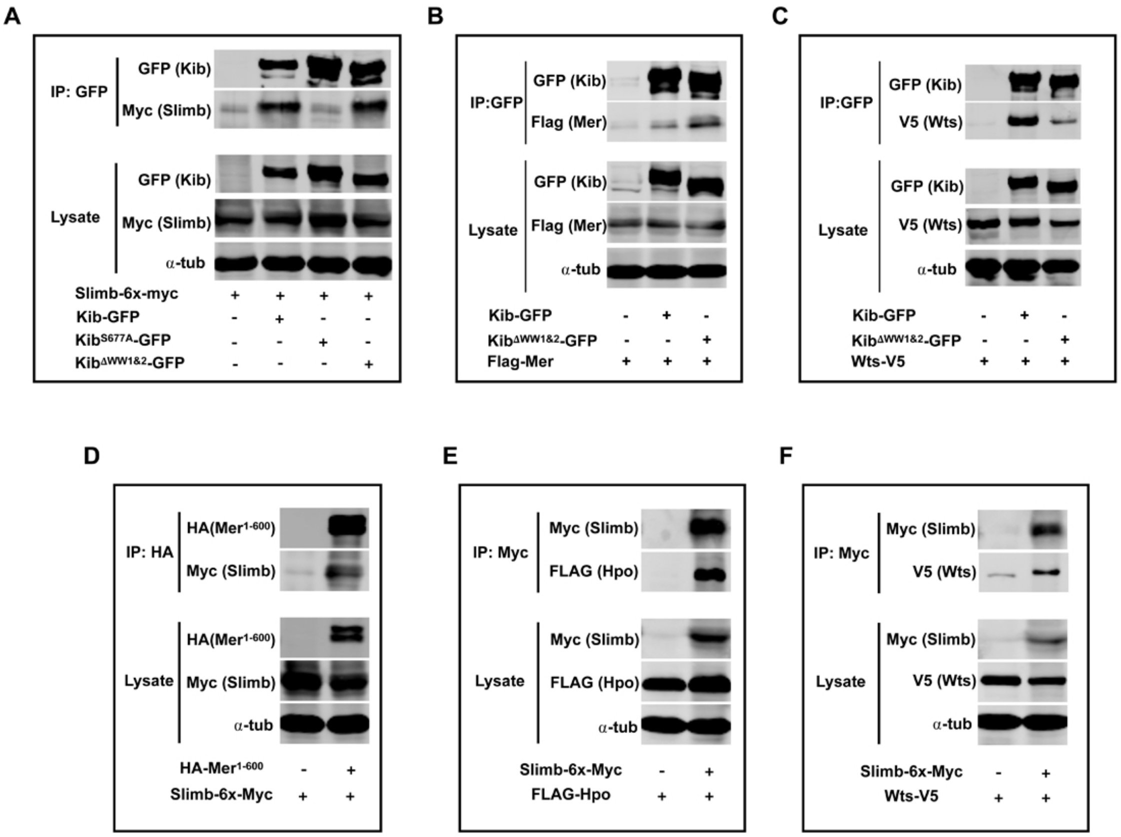

**Figure S6, related to Figure 6.**
**Complex formation and Kib degradation.**

A-C) Co-IP of wild-type Kib or $Kib^{\Delta WW1\&2}$ with Slimb (A), Mer (B), or Wts (C).
D-F) Slimb forms a complex with $Mer^{1\text{-}600}$ (D), Hpo (E), and Wts (F).
All experiments were performed using lysates from transfected S2 cells.

### Supplemental Figure S7

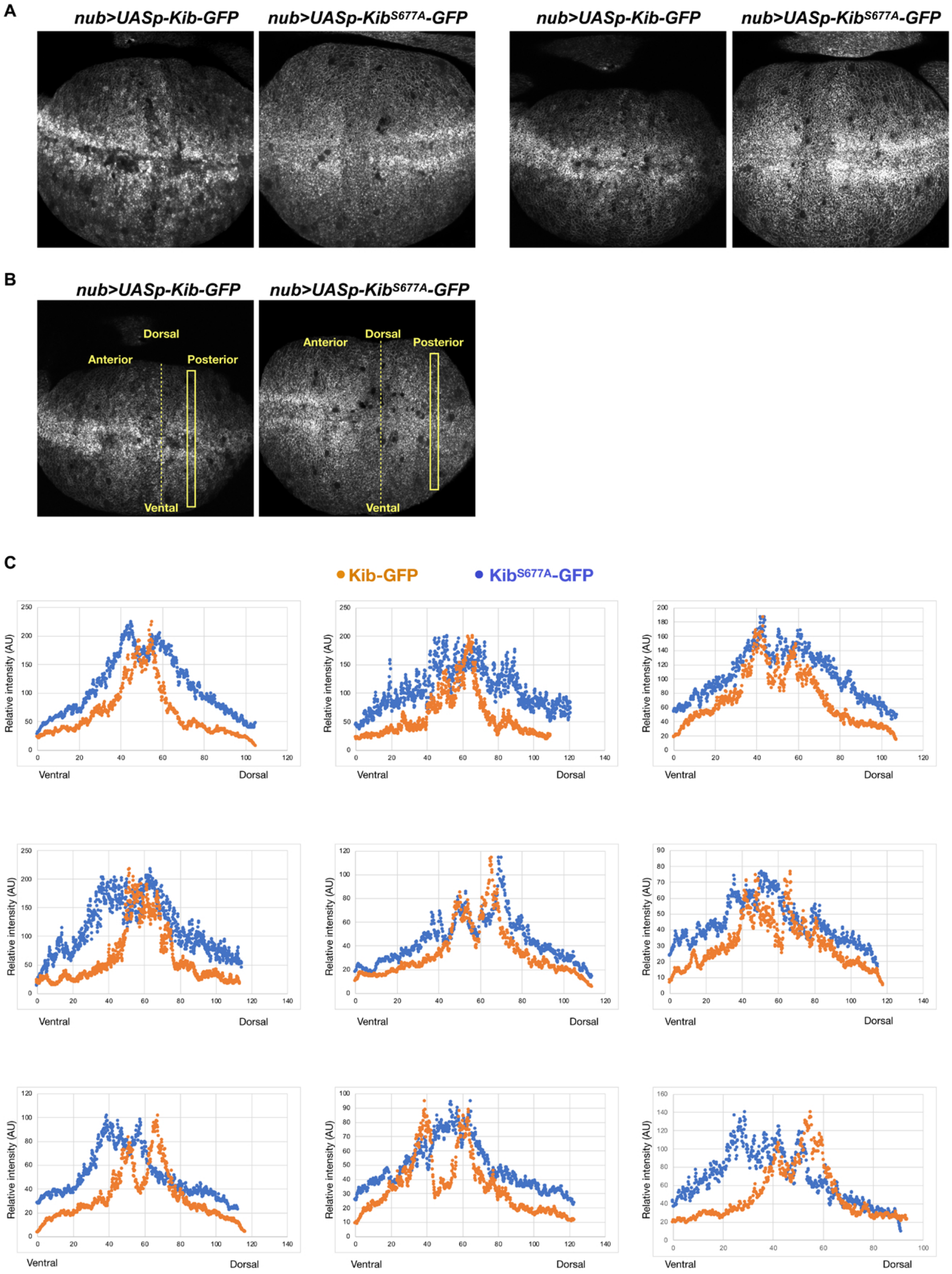

**Figure S7, related to Figure 7.**

**Kib degradation is not uniform across the pouch region of the wing imaginal disc.**

A) Additional examples of UASp-Kib-GFP and UASp-Kib$^{S677A}$-GFP expression patterns under the *nub>Gal4* driver focused on the wing pouch, which produces the adult wing blade.

B) To generate the intensity profiles in (C), rectangular selections of equal length were drawn on each disc either in the anterior or posterior region (same side was used for each pair). Intensity values from UASp-Kib-GFP discs were normalized to the maximum value of UASp-Kib$^{S677A}$-GFP discs.

C) Intensity profiles of 9 pairs of wing imaginal discs expressing UASp-Kib-GFP or UASp-Kib$^{S677A}$-GFP under the *nub>Gal4* driver. In each case, wild-type Kib intensity drops more severely from the distal (middle) to proximal (most dorsal or ventral) part of the wing pouch.
