## Supplemental Methods for "Yorkie-independent negative feedback couples Hippo pathway activation with Kibra degradation"

### Supplemental Materials and Methods

### Table S1: Antibodies

| Antibodies | Source | Identifier |
|---|---|---|
| Guinea pig anti-Ex | (Maitra et al., 2006) | RRID:AB_2568722 |
| Mouse anti-FLAG | Sigma | Cat#F1804; RRID:AB_262044 |
| Rat anti-Ecad | DSHB | Cat#AB_528120; RRID:AB_528120 |
| Guinea pig anti-Sd | (Guss et al., 2013) | RRID:AB2567874 |
| Guinea pig anti-GFP | This paper | NA |
| Rabbit anti-GFP | Michael Glotzer | NA |
| Mouse anti-Hpo | (Su et al., 2017) | NA |
| Rabbit anti-HA | Santa Cruz | Cat#sc-805; RRID:AB_631618 |
| Mouse anti-Myc (9B11) | Cell Signaling | Product #2276 |
| Mouse anti-V5 | GenScript | Cat# A01724-100 |
| Mouse anti-alpha tubulin | Sigma | Cat# T 9026 |

### Table S2: Fly stocks

| Stocks used: | Source |
|---|---|
| *Kib::GFP* | (Su et al., 2017) |
| *$w^{1118}$; $yki^{B5}$ {yki-YFP}* | This study |
| *Mer4 19AFRT/FM7actGFP; Sco/CyodfdYFP* | This study |
| *Exe1 40AFRT/CyodfdYFP; MKRS/TM6, Tb* | This study |
| *19AFRT $sd^{47}$M* | (Wu et al., 2008) |
| *hpoBF33 42DFRT* | (Jia, 2003) |
| *ban3-GFP* | (Matakatsu and Blair, 2012) |
| *UAS-Mer RNAi (II)* | This study |
| *UAS-Sav RNAi (II)* | BL28006 |
| *UAS-Hpo RNAi* | VDRC 104169 |
| *UAS-Wts RNAi* | VDRC 106174 |
| *UAS-Ex RNAi* | VDRC 109281 |
| *UAS-Crb RNAi* | VDRC 39177 |
| *UAS-Yki RNAi (III)* | VDRC 40497 |
| *UAS-Slimb RNAi* | BL33898 |
| *UAS-Cul1 RNAi* | BL29520 |
| *UAS-SkpA RNAi* | BL32870 |
| *UAS-Mahj RNAi* | BL 34912 |
| *UAS-Nedd4 RNAi* | BL 34741 |
| *UAS-POSH RNAi* | BL 64569 |
| *UAS-POSH* | BL 58990 |
| *UAS-Su(dx) RNAi* | BL 67012 |
| *UAS-Herc4* | (Aerne et al., 2015) |
| *UAS-Smurf RNAi* | BL 40905 |
| *UAS-Fbxl7 RNAi* | VDRC108628 |
| *UAS-Ed RNAi* | BL 38243 |

| | |
|---|---|
| *UAS-Ft RNAi* | BL 34970 |
| *UAS-Dachs-V5* | (Mao, 2006) |
| *UAS-Tao1 RNAi* | VDRC 17432 |
| *UAS-Mats RNAi* | BL 34959 |
| *UAS-Pez RNAi* | BL 33918 |
| *Ey>Flp 19AFRT Ubi-GFP; Ubi-RFP 42DFRT* | (Koontz et al., 2013) |
| *Ubi-Kib-GFP 86Fb* | This study |
| *UASp-Kib-GFP-FLAG 86Fb* | This study |
| *UASp-Kib$^{S677A}$-GFP-FLAG 86Fb (this study)* | This study |
| *Ubi-Kib-GFP-FLAG VK37* | This study |
| *Ubi-KibΔWW1-GFP-FLAG VK37* | This study |
| *Ubi-KibΔWW2-GFP-FLAG VK37* | This study |
| *Ubi-KibΔWW1&2-GFP-FLAG VK37* | This study |
| *Ubi-Kib1-857-GFP-FLAG VK37* | This study |
| *Ubi-Kib484-1288-GFP-FLAG VK37* | This study |
| *Ubi-Kib858-1288-GFP-FLAG VK37* | This study |
| *Ubi-KibΔCC1-GFP-FLAG VK37* | This study |
| *Ubi-KibΔCC2-GFP-FLAG VK37* | This study |

### Table S3: Primers

| Primers | Sequence |
|---|---|
| Gibson to pUbi_For | TTCTTCCCGCAGATAATCCAAATCGTTAACAGATCTGCGG |
| Gibson to pUbi_Rev | AAGTAAGGTTCCTTCACAAAGATCC |
| Gibson to pMT_For | TCAGTGCAACTAAAGGGAATTCGATATCTCGTTAACAGATCTGCGG |
| Gibson to pMT_Rev | AGGTCGACTCTAGAGGATCCCCGGGAAAGATCCTCTAGAGGTTACTTG |
| ΔWW1 For | AGCAACAACACCACAGCGACTGCTACACAAAGCCGCAGACTTT |
| ΔWW Rev | CTTTGTGTAGCAGTCGCTGTGGTGTTGTTGCT |
| ΔWW2 For | GACTTTCGAGGATTGTGTGGGCGAGTGGAAGACTGTCCAGGAGCA |
| ΔWW2 Rev | TCTTCCACTCGCCCACACAATCCTCGAAAGTCT |
| ΔWW 1&2 For | AGCAACAACACCACAGCGACGAGTGGAAGACTGTCCAGGAGCA |
| ΔWW 1&2 Rev | TCTTCCACTCGTCGCTGTGGTGTTGTTGCT |
| ΔCC1 For | TCGATGAGTCGCCACGATCCGTACACGGAACGGGGCATGAACA |
| ΔCC1 Rev | GTTCCGTGTACGGATCGTGGCGACTCATCGAA |
| ΔCC2 For | ACCTGAACGGAGGAGCCCGTTTCTCGGAGAGCACCTTCTCCATTAGCAGT |
| ΔCC2 Rev | TGCTCTCCGAGAAACGGGCTCCTCCGTTCAGGT |
| 484-1288 For | TTCGTTAACAGATCTGCGGCCGCGCCACCATGAGTAAGAGCGCCTTGAGCTTCAC |
| 484-1288 Rev | TCGCCCTTGCTCACGCCGGAGCCGGTACCGGACACCTCCACGCCGTAGTTGCGA |
| 1-857 For | TTCGTTAACAGATCTGCGGCCGCGCCACCATGCCGAATCTGCAACAAACCGC |
| 1-857 Rev | TCGCCCTTGCTCACGCCGGAGCCGGTACCGGACTCATCCGACGACTCCTCCCGGTTG |
| 858-1288 For | TTCGTTAACAGATCTGCGGCCGCGCCACCATGTCCACCATTACATCCTCCCAGAC |
| 858-1288 Rev | TCGCCCTTGCTCACGCCGGAGCCGGTACCGGACACCTCCACGCCGTAGTTGCGA |
| KibS677A For | CCGGCGATGCTGGCGTCTTCGAG |
| KibS677A Rev | CCACGGATTCATTGCTGACCGC |
